# Unveiling epigenetic regulatory elements associated with breast cancer development

**DOI:** 10.1101/2024.11.12.623187

**Authors:** Marta Jardanowska-Kotuniak, Michał Dramiński, Michał Własnowolski, Marcin Łapiński, Kaustav Sengupta, Abhishek Agarwal, Adam Filip, Nimisha Ghosh, Vera Pancaldi, Marcin Grynberg, Indrajit Saha, Dariusz Plewczynski, Michał J. Dąbrowski

## Abstract

Breast cancer is the most common cancer in women and the 2nd most common cancer worldwide, yearly impacting over 2 million females and causing 650 thousand deaths. It has been widely studied, but its epigenetic variation is not entirely unveiled. We aimed to identify epigenetic mechanisms impacting the expression of breast cancer related genes to detect new potential biomarkers and therapeutic targets. We considered The Cancer Genome Atlas database with over 800 samples and several omics datasets such as mRNA, miRNA, DNA methylation, which we used to select 2701 features that were statistically significant to differ between cancer and control samples using the Monte Carlo Feature Selection and Interdependency Discovery algorithm, from an initial total of 417,486. Their biological impact on cancerogenesis was confirmed using: statistical analysis, natural language processing, linear and machine learning models as well as: transcription factors identification, drugs and 3D chromatin structure analyses. Classification of cancer vs control samples on the selected features returned high classification weighted Accuracy from 0.91 to 0.98 depending on feature-type: mRNA, miRNA, DNA methylation, and classification algorithm. In general, cancer samples showed lower expression of differentially expressed genes and increased *β*-values of differentially methylated sites. We identified mRNAs whose expression is well explained by miRNA expression and differentially methylated sites *β*-values. We recognized differentially methylated sites possibly affecting NRF1 and MXI1 transcription factors binding, causing a disturbance in *NKAPL* and *PITX1* expression, respectively. Our 3D models showed more loosely packed chromatin in cancer. This study successfully points out numerous possible regulatory dependencies.

## Introduction

In 2021, the WHO announced that for the first time in 20 years the most commonly diagnosed cancer in the world was not lung cancer but breast cancer. According to the GLOBOCAN report publishing cancer statistics for 2020 based on data from 185 countries and 36 different types of cancer, 2.3 million people were affected by breast cancer, and 684,996 people died from it [1]. It means that currently almost one in four oncology female patients develops breast cancer. Early stage cancer detection is crucial to apply the most effective therapy available [2]. Due to the heterogeneous nature of breast cancer and the large amount of information to be considered, the implementation of appropriate treatment is extremely difficult [3].

The effectiveness of cancer therapies is closely related to diagnostic accuracy. Inclusion of molecular features in the classification of cancers [4,5] has allowed for the development of targeted treatment. Further molecular studies aimed at detecting cancer markers and potential drug targets are essential to further improve the available therapies. Large molecular databases, which are extremely rich in information, but also burdened with a certain level of information noise, which is a significant analytical challenge, especially when the number of samples available is limited and data dimension is high. In such a case, a risk of obtaining false positives and false negatives may be higher, because from the statistical perspective the problem is ill-defined [6]. To overcome this challenge, in this research we applied the Monte Carlo Feature Selection and Interdependency Discovery (MCFS-ID) algorithm [7] to reveal significant signals related to breast cancer in various molecular datasets [8]. The MCFS-ID has been successfully used in a broad range of scientific disciplines, including oncology, virology and cardiology [9–13].

It is a known fact that breast cancer is associated with multiple DNA mutations and genome rearrangements that affect cell physiology, resulting in gene expression changes [14,15]. Yet, it has been shown that DNA alterations alone cannot fully explain breast cancer development, and in recent years there has been accelerated research toward the epigenetic regulation of cancer-related gene expression [16–20].

One of the most important and well-studied epigenetic modifications is DNA methylation. Methylation of gene promoters and regulatory regions plays an essential role in regulating gene expression and shows high variation across cell types. Its deregulation is associated with tumorigenesis [21–23] and was demonstrated to have a role in predicting patient survival [17]. The DNA binding affinity of multiple transcription factors (TFs) relies on DNA methylation patterns [24,25]. Interestingly, DNA hypo-methylation is present in the regulatory regions of oncogenes promoting tumorigenesis [26–28], while hyper-methylation is frequently connected with the silencing of tumor suppressor genes [29–31], but other research shows that these patterns may be more complex and depend on genomic location of the methylation alterations [32,33]. That is why a further large-scale analysis of locus-specific DNA methylation patterns in relation to TF affinity and the level of gene expression may bring novel knowledge about cancer biomarkers and deregulated biological pathways that promote tumorigenesis.

Similarly to DNA methylation, miRNAs can also act as epigenetic regulators [34] that may down-regulate gene expression of several genes in breast cancer. Additionally, miRNAs may regulate gene expression by inducing mRNA turnover and thus silencing protein-coding mRNAs [35] and influencing the development and drug resistance of breast cancer [36,37]. Another epigenetic mechanism known to contribute to cancerogenesis is 3D chromatin structure alterations, which can be affected by several molecular elements including point mutations [38] or DNA methylation levels [39]. Such alterations were shown to disrupt gene expression in many cancers [40,41]. Therefore, defining the distances between regulatory elements and their target promoters in 3D chromatin structure may provide insights into the underlying mechanisms of genomic regulation also in breast cancer.

In the present study we applied the MCFS-ID algorithm to extract the significant transcriptomic and DNA methylation features from The Cancer Genome Atlas (TCGA) dataset that could distinguish between healthy and cancerous tissues. Subsequently, we conducted analyses of mRNA expression, DNA methylation, detection of TF motifs, miRNA potential targeting by drugs and modeling of 3D chromatin structure. This integrative approach helped to reveal the biological importance of the selected features, as well as the direct and indirect connections between them and their impact on the initiation and development of breast cancer.

## Materials and Methods

### Data collection

This study is based on breast cancer data obtained from The Cancer Genome Atlas (TCGA) including: mRNA, miRNA expression and DNA methylation levels [https://www.cancer.gov/tcga]. Data was filtered as follows: (1) all attributes having zero variance across samples were removed; (2) only female samples were included in the study. The final dataset consisted of 1191 samples taken from 1068 female patients (123 patients donated both normal and cancerous tissues). Out of 1191 samples only 381 were complete among mRNA, DNA methylation and miRNA data. They consisted of 328 cancerous and 53 normal samples (Table 1 and Fig 1). The remaining set of samples (incomplete among all datasets) were used as testing sets in separated classification experiments described in sections “Detection of significant features using MCFS-ID algorithm” and “Detection of potential breast cancer biomarkers using the MCFS-ID algorithm”.

**Fig 1.**
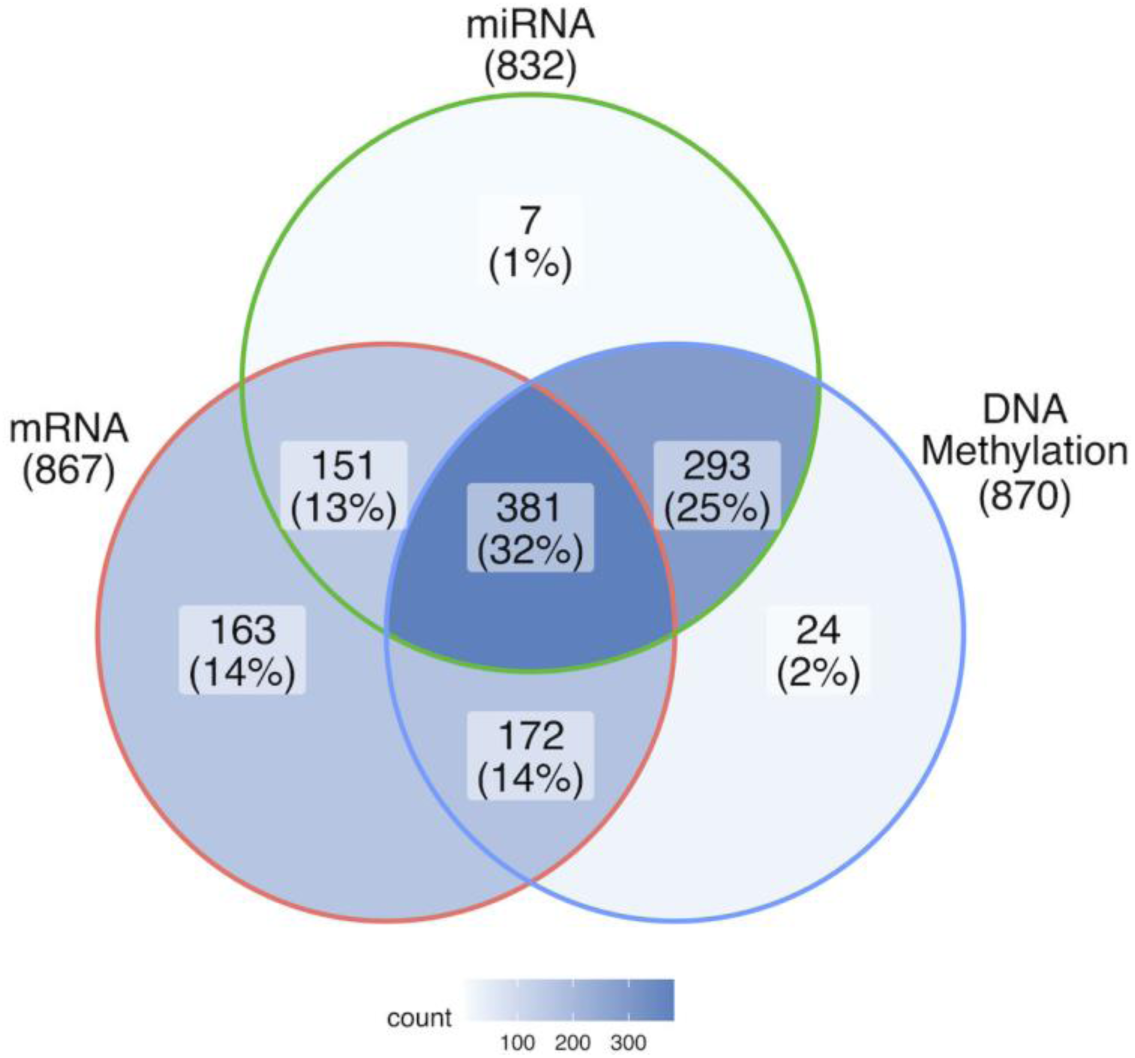
Number of samples with the complete data for a given dataset type. The overlap of 381 samples that contain the complete data for all three dataset types was used in the feature selection procedure.

**Table 1.** Input data description.

| Data Type | Unit of measurement | Number of total samples | Number of normal samples | Number of features |
| --- | --- | --- | --- | --- |
| mRNA expression | reads per kilobase million | 867 | 99 | 20524 |
| DNA methylation | beta-value | 870 | 97 | 396065 |
| miRNA expressions | reads per million miRNA mapped | 832 | 86 | 897 |

### Detection of significant features using MCFS-ID algorithm

Our analysis utilized the Monte Carlo Feature Selection and Interdependencies Discovery (MCFS-ID). This algorithm allows the user to perform a supervised feature selection introduced in [42]. MCFS-ID generates a ranking of features based on their potential to distinguish records between classes, e.g., cancerous vs normal. It also enables the prediction of continuous values and allows the user to discover possible interdependencies between features.

The algorithm builds thousands of decision (or regression) trees on randomly selected subsets of data samples and attributes. The relative importance (RI) score for each feature is calculated based on all decision trees and nodes built on that feature: the number of samples split by the node, information gain of the node and the predictive quality of the trees using that feature. The RI score is used to build the ranking of all input features. The ranking signifies which attributes are best to be used in classification or regression tasks. Additionally the algorithm provides a RI cutoff that assures that attributes that exceed it are better for predictions than attributes with random values, which might distinguish classes by pure chance. The upper part of the feature ranking cut off by the RI value constitutes the significant features set.

The Interdependency Discovery function of the MCFS-ID algorithm allows the user to find links between features that amplify each other in the classification task. Decision or regression trees used for calculation of RI score are also used to generate feature interdependency scores. In this case the score for the pair of features (parent/child nodes in the tree) is generated using information gain of the child node multiplied by its associated number of samples expressed as a fraction of samples in the parent node.

The resulting scores indicate which features amplify the prediction powers of other features. The result of this algorithm has the form of a directed graph where features are visualized as nodes, and the thickness of edges symbolizes the strength of this amplification. For better clarity, and to avoid false positives, MCFS-ID allows us to cut off edges that are weaker than those which might be caused by random patterns occurring in the data. Interdependency graphs were generated using the *build.idgraph* function from the *rmcfs* package. For more details of the MCFS-ID, see [7].

To establish a final list of significant features for each data type, four runs of the MCFS-ID algorithm were performed (Fig 2).

**Fig 2.**
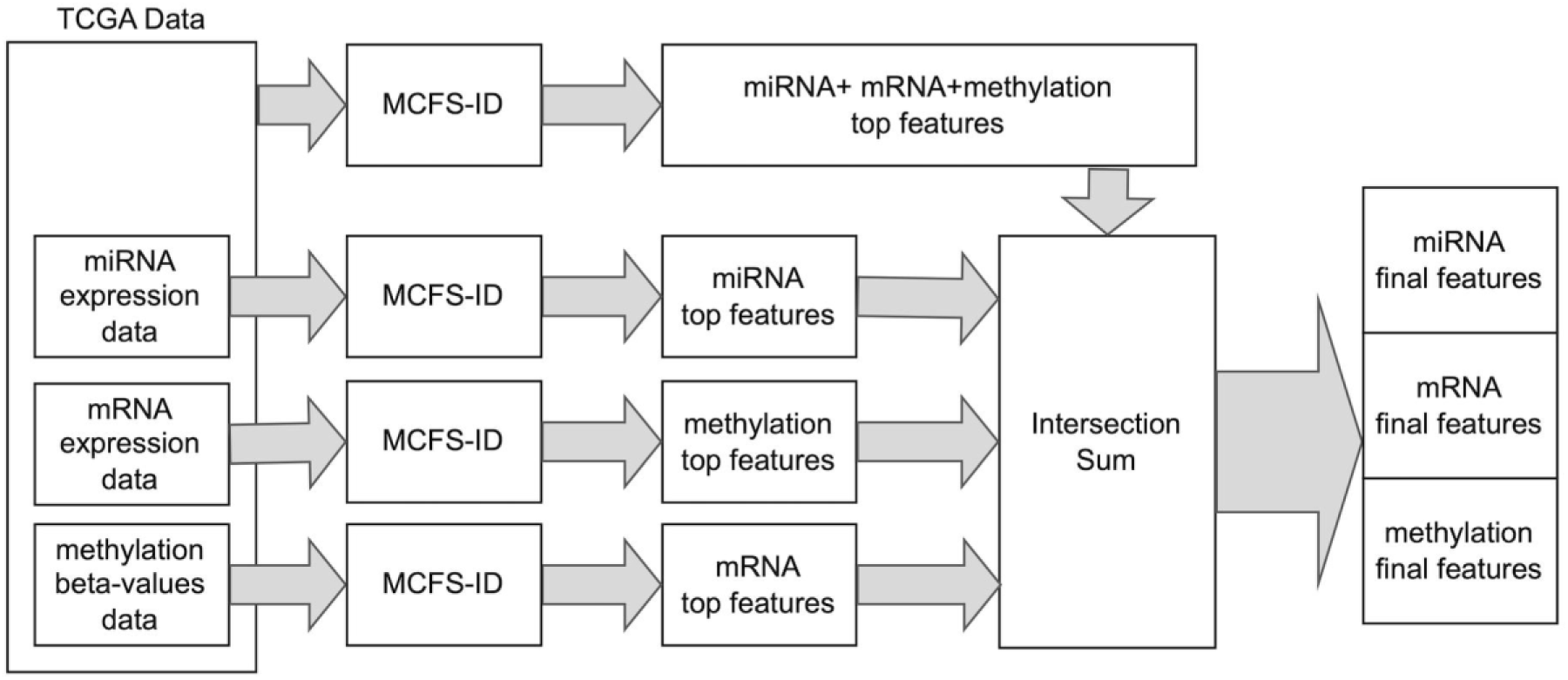
A flowchart describing selection of features identified as significant in cancer prediction, including attributes that are significant within one separated type of data and significant when combining all types of data together. This procedure will be called the main MCFS-ID experiment later in the paper and features selected by the algorithm as the significant set of mRNA/miRNA genes or DNA methylation features.

First of all, a selection of features was performed on the full data set consisting of mRNA expression, DNA methylation and miRNA expression in 381 samples. This approach allowed for capturing the relationships between attributes from different types of data sets. Next, a feature selection was performed on the same dataset set but for each data type separately (Fig 2), where the sample size differed depending on a data type (Table 1, Fig 1). In all of these four runs the features were selected by their ability to differentiate between normal and cancerous samples. Running MCFS-ID on data of all types allowed us to find features significant in conjunction with features of other types. Attributes derived from this run are to be called significant along with all categories. Separate runs for each data type capture these less important (but still important) features that would be under the relevance threshold of MCFS-ID algorithm in case they were part of a larger set of features. The analysis was conducted using version 1.3.1 of the *rmcfs* package from the CRAN repository in R 3.6.3. The default parameters were used except: splitSetSize=200; mode=2; cutoffPermutations=20.

Additionally, to validate the quality of selected important features, two different classification models: SVM (support vector machines) and RF (random forest) were trained on the 381 complete samples and tested later on the test samples that were not used in the feature selection phase. For each type of the data there are over 400 unique test-ready samples that do not contain values for all categories and were used to obtain weighted Accuracy of classification (wAcc) for each category separately.

### Descriptive analysis of significant mRNA genes

To annotate genes as down- or over-expressed in cancer samples, the log2 fold change was calculated for each of the significant genes (returned by MCFS-ID). Distribution of gene expression was verified with the Shapiro-Wilk test and depending on the obtained results the Wilcoxon test was applied. Next, functional annotation of significant genes was performed using the Enrichr web server [43] followed by the Benjamini–Hochberg correction for multiple testing. Initially the q-value was set to 0.05 but because of a huge number of significant terms returned, in one (over-expressed mRNA) analysis the threshold was set to 0.001. Moreover, based on the set of significant genes descriptions, gathered from various molecular databases provided by BioMart (such as NCBI, Gene Ontology, KEGG, Reactome, WikiPathways, Biocarta), Natural Language Processing (NLP) methods were applied to group these genes into functionally/descriptively similar clusters and retrieve sets of key words to describe each of the returned cluster. To perform this operation, each gene was represented as the text document (combined from all available descriptions), and, later, used to calculate TF-IDF (term frequency–inverse document frequency), as a bag of words/terms. When each gene is represented by a TF-IDF vector it is possible to calculate cosine similarity between the documents and a hierarchical clustering model can be built based on these similarities. Finally, discovering a set of unique keywords that characterize each cluster provides a good functional biological overview of input genes and groups that should be studied more closely.

### Descriptive analysis of significant DNA methylation sites

DNA methylation sites selected by MCFS-ID (hereafter differentially methylated sites, DMSs) were mapped to the specific genomic regions. Original DMSs positions were converted from hg19 to hg38 using LiftOver [44]. Intersections of genomic regions i.e. CpG islands (CpGIs) or promoters with DMSs were done using bedtools (v2.29.2) [45]. The locations of CpG islands (CpGI) were taken from the UCSC database represented in hg38 genome assembly (https://hgdownload.cse.ucsc.edu/goldenpath/hg38/database/cpgIslandExt.txt.gz). The shores were defined as the regions flanking the CpGI by 2000 bp up- and down-stream, the shelves as the regions flanking the shores by +/-2000 bp, and the open seas as the regions between the shelves. Next, to prepare the background distribution of cytosines across these four region types, all cytosines, which are included in the Human Methylation 450K BeadChip (Illumina 450K) were mapped to CpGIs, shores, shelves, and open sea regions. Sites covered by the Illumina 450K represented the background distribution of cytosines across the named genomic regions and allowed to verify (chi-squared test) whether DMSs show any specific distribution. Afterwards, the distribution of *β*-values of cancer vs normal samples was compared separately for CpGIs, shores, shelves and open sea regions using the Wilcoxon test. Just to note, that *β*-values represent the ratio of the intensity of the methylated bead type to the combined intensity of a locus and obtains values between 0 and 1 [46]. Finally, we tested whether the distributions of hypo-, medium- and hyper-methylated cytosines in cancer samples differed across the four region types (chi-squared test). Bonferroni correction was applied for multiple testing corrections for the aforementioned analysis.

To annotate DMSs to promoters or gene bodies, the gene positions from Ensembl (https://www.ensembl.org/Homo_sapiens/Info/Index) were taken and their promoters were set +/-2000 bp around the TSS. DMSs assigned to the promoter or the gene body established a pair: a DMS and its target gene. DMSs within intergenic regions were paired with their target genes using bedtools *closest* function. Assignment of hyper-, hypo-, medium-methylated *β*-values was performed on the basis of the log2 fold change (log2FC) between sample types (cancer vs normal). Sites with log2FC ≤ -1 were labeled as hyper-methylated in cancer; log2FC ≥ 1 as hypo-methylated DMSs in cancer; the remaining sites were labeled as medium-methylated. To evaluate the significance of the observed log2FC, first the Shapiro-Wilk test was used to verify whether the data was normally distributed and, based on the test results, a nonparametric Wilcoxon test with FDR correction was chosen to verify the null hypothesis: that there is no difference in distribution of *β*-values for DMSs between cancer and normal samples.

To detect putative regulatory regions, the association between gene expression and *β*-value of DMSs located 1Mb upstream/downstream from that gene TSS was verified by calculation of the Spearman correlation with FDR correction, defining significant correlations when Spearman’s |rho| ≥ 0.6 and FDR ≤ 0.05. Moreover, to assign DMSs to specific chromatin states, we used chromatin state annotations for the MCF-7 breast cancer cell line (GSE57498). The data were converted from original annotation Human GRCh37/hg19 to GRCh38/hg38 annotation using LiftOver [44]. Positions indicating acetylation of the lysine 27 of the histone H3 protein (H3K27ac) for MCF-7 cell line were taken from the ENCODE database (https://www.encodeproject.org/files/ENCFF621API/). The sites included in the Illumina 450K panel were intersected with the ranges of chromatin states to achieve the general distribution of cytosines across chromatin states. Next, from the significant 2006 DMSs, hyper- and hypo-methylated DMSs were selected and independently intersected with chromatin states to unveil their distribution across chromatin states. To verify whether the obtained distributions were specific, 2006 random loci were drawn 1000 times from the Illumina 450K panel. Each time a set of those drawn loci was intersected with the chromatin state ranges to compute the percentage of loci assigned to specific chromatin state. Next, the logarithm of fold change between percentage of hyper-methylated DMSs and mean percentage of randomly drawn loci for each chromatin state was computed. The same procedure was applied to hypo-methylated sites, generating empirical p-values of significance for these overlaps.

Next, to evaluate the impact of DMSs on the patients’ survival, a multivariate log rank test (*lifelines.statistics.multivariate_logrank_test function in Python*) was used. Samples were split into high and low methylated groups defined by a median of methylation level. Additionally, to confirm that the number of DMSs, discovered to have a significant impact on survival, differs significantly from the number of such sites selected randomly, a bootstrapping technique (sampling 100 times) was applied. After each sampling of 2006 random sites from all sites present in Illumina 450K panel, their impact on survival was tested and the number of sites with p-value below 0.05 was noticed. Next, we counted how many times the number of significant sites (having impact on survival) found in a random set was greater than the number of significant sites (having impact on survival) obtained for 2006 DMSs.

### Descriptive analysis of significant miRNA genes

To study the suppressive impact of significant miRNAs (those returned in the main MCFS-ID experiment) on mRNA gene expression, the selection of over-expressed miRNAs genes in cancer was performed based on log2FC. The genes were defined to be over-expressed in cancer if log2FC ≤ -0.1 (S1A Fig). Next, for the selected miRNAs over-expressed in cancer, their mRNA target genes were assigned using the MicroRNA Target Prediction Database [47] and Spearman correlation was calculated for the obtained miRNA-mRNA pairs. The correlations where *rho* ≤ -0.2 and adjusted p-value ≤ 0.05 were considered in further analysis. Next, for the mRNA genes that significantly correlated with miRNA genes using the STRING database [48], the protein-protein interaction identification was performed and based on the number of interactions among proteins, the top 50 proteins were selected. Next using KEGG pathways and gene ontology biological processes (GO BP) the enrichment analysis on those top 50 proteins was performed. Additionally, to discover putative drugs associated with over-expressed miRNAs, the Enrichr Database was searched with the same set of proteins.

Finally, to characterize the set of significant miRNAs returned by MCFS-ID, we searched them in the miR+Pathway database [49] that contains information about mRNA-miRNA connections and 150 KEGG pathways linked with mRNAs. The mRNAs linked with the searched miRNAs were intersected with 590 mRNAs returned in the MCFS-ID experiment. Biological functions of the resulting mRNAs were verified on NLP cluster key words (described in section “Descriptive analysis of significant mRNA genes”)

### Detection of significant miRNA and methylations in the context of predicting mRNA expression levels

To discover more complex interactions between the top 590 most significant mRNA genes (obtained from the main MCFS-ID experiment described in section “Detection of significant features using MCFS-ID algorithm”) and two other types of molecular data (DNA methylation and miRNA expression) two additional sets of MCFS-ID experiments were performed (Fig 3). Both sets of experiments were based on the same idea of running MCFS-ID algorithm on all top significant 590 mRNA features obtained from the main MCFS-ID runs. Each of those mRNA features was used as a target variable and miRNAs or DNA methylation were used as predictor features. Thus finally two different feature rankings were obtained. In the case of DNA methylation, for each run on a different target mRNA gene, methylation loci within the chromosome of the gene were explored. This limitation heavily reduced the calculation time - the number of all DNA methylation sites is almost 400k and it is biologically justified to focus on such relations within one chromosome [50]. In both sets of experiments, final cross validation of the result was based on a regression tree modeling and calculation of Pearson correlation between predicted value and observed mRNA gene expression level.

**Fig 3.**
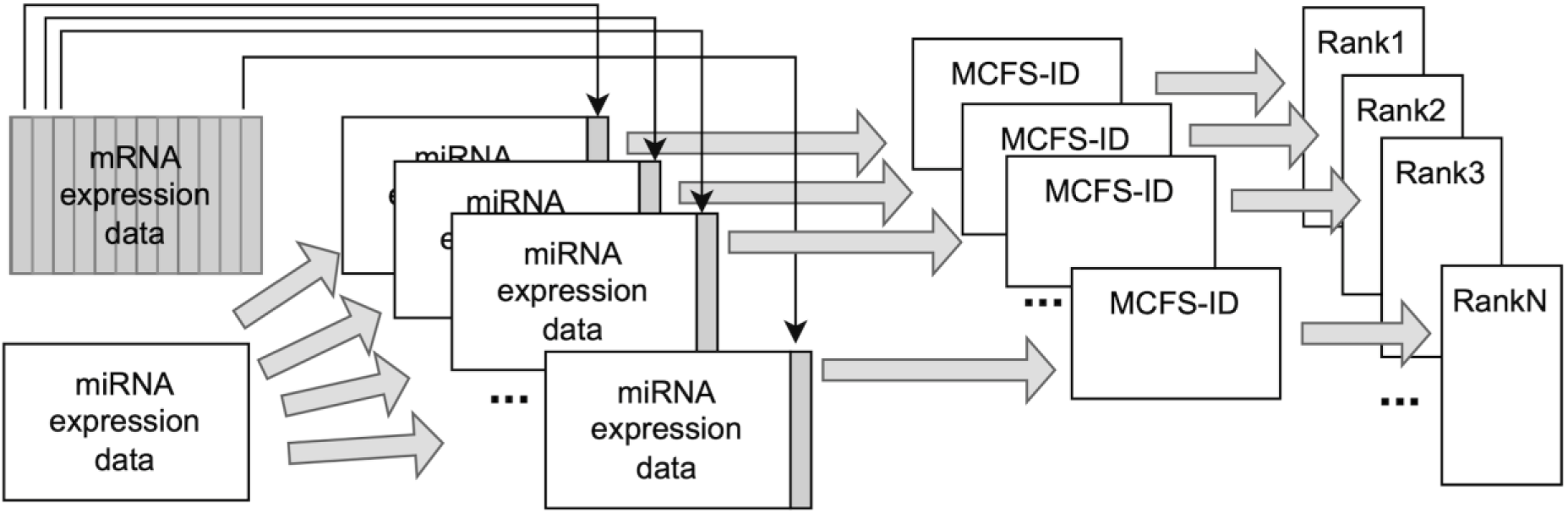
A flowchart describing selection of miRNA genes found to be significant in prediction of mRNA gene expression levels. For each significant mRNA (selected by the main experiment described in section “Detection of significant features using MCFS-ID algorithm”) treated as a target variable, a separate MCFS-ID experiment was performed. The same procedure was used for DNA methylation as predictors instead of miRNA expression levels.

### Descriptive analysis of associations between DMS and TFs

To predict transcription factors (TFs) whose binding affinity to DNA sequence may be changed due to differential DNA methylation, the sequences surrounding each DMS (+/- 20 bp) were generated using bedtools getfasta function (v2.29.2) [45]. To identify TF motifs, the HOCOMOCO database of position weight matrices (PWMs) [51] together with the PWMEnrich tool [52] were used with the following settings: (i) sequences background build based on randomly selected fasta sequences and (ii) motif significance cutoff p-value ≤ 0.001. Next, to detect the exact Transcription Factor Binding Site (TFBS) positions of the motifs that passed the threshold, the online FIMO tool [53] from the MEME Suite 5.0.5 was applied with significance threshold p ≤ 0.0001. The returned TFBS were intersected with DMS to keep only these TFBS within which any DMS was confirmed, to ensure that differential DNA methylation may affect binding affinity. To group TF motifs, returned by PWMEnrich, based on the heterogeneity of their PWMs, the STAMP tool [54] was used with Pearson Correlation Coefficient as a measure of distance. Sequence alignment was performed using an ungapped Smith-Waterman algorithm with the iterative refinement multiple alignment strategy. Visualization of the clustering results was performed with UPGMA [54]. Further, functional annotation of genes encoding TFs was done using the Enrichr web server [43] with the Benjamini–Hochberg correction for multiple testing and significance threshold q-value ≤ 0.05.

### Models of regulatory networks

To investigate whether the putative associations between TFs and DMSs within binding sites of these TFs have similar patterns in cancer vs normal samples, the additional analysis using the MCFS-ID algorithm was performed. The input decision table consisted of 484 patients for whom both mRNA and DNA methylation datasets were available. Features, namely TFs coding genes and DMS within DNA sequences of motifs detected for that TFs were analyzed. The returned significant features, together with cancer/normal tissue type, were used as explanatory variables to train a set of linear models to predict mRNA expression of a target gene. Each time a different mRNA gene (target gene), whose promoter overlapped with a set of explanatory variables (TFs and DMS), was used as a dependent variable to build a single linear model. Finally, for the best fitted models (adjusted p≤0.05, R^2^ > 0.5), the feasible biological relationships between DMS, TFs and their target genes were visualized. These target genes were also used to discover direct and the closest indirect associations between them and other genes by the literature systematic review and interaction graphs obtained through the Pathway Commons online tool [55]. Based on the associations found, a final graph of connections between identified target genes and other genes was created and visualized.

### The visualization of chromatin 3D structure of selected *loci*

In order to visualize the putative functional association between genes, DNA methylations significant in breast cancer, and enhancer regions, the chromatin structures of the *FXYD1* and *NKAPL loci* were generated using 3D-GNOME [56] and Spring Model (SM) [57] polymer simulation methods.

3D-GNOME is a chromatin 3D structure modeling method that uses a multiscale bead-on-a-string approach and a Monte Carlo simulated annealing algorithm [56]. It models chromatin interactions mediated by specific proteins, based on high-frequency PET (multiple paired end tags mapped on two genomic loci) cluster interactions and singletons (single paired end tag). The algorithm uses a tree structure to manage the relationships between different levels of genomic organization (chromosomes, segments containing a topological domain, and chromatin interaction anchors), simulating their spatial positions independently by minimizing an energy function based on high-frequency chromatin PET cluster interactions and energy terms. Next, sub-anchor beads are added between neighboring anchor beads to model chromatin loops, and their positions are again simulated by minimizing energy. Finally, the algorithm refines the loop shape using a singleton interaction heatmap and motif orientation. The 3D-GNOME models were generated based on cohesin mediated Chromatin Interaction Analysis by Paired-End Tag sequencing (ChIA-PET) data for the hTERT-HME1 (normal) and MCF-7 (cancer) cell lines (ENCODE Accession ID: ENCSR991JXX - hTERT-HME1, ENCSR255XYX - MCF-7) [58].

The Spring Model represents polymers as a collection of points in three-dimensional space using the beads-on-chain approach. In the resolution chosen by the user, each bead represents a segment of DNA of the same length. In this study, the chromatin models with a resolution of 1 kbp were constructed, where each bead represents 1,000 base pairs. If there was a spatial interaction between beads in the polymer model, harmonic bonds were used to connect pairs of interacting beads by springs. The spring-based pairwise forces are subjected to energy minimization in a SM polymer simulation. In order to establish the final 3D structure of the polymer fiber with the set of experimentally identified contacts, the SM simulation undertakes the global energy minimization given the data-driven forces represented by the springs and polymer chain parameters (such as stiffness). The initial conformation of the polymer was given as a circular 3D structure of polymer fiber. 3D models generated by the Spring Model approach were built using Promoter Capture Hi-C (PCHi-C) [59] data for MCF-10A (healthy) and MCF-7 (cancer) breast tissues to get a promoter centric view.

## Results

### Detection of potential breast cancer biomarkers using the MCFS-ID algorithm

Our study aimed to verify if there are significant molecular features and interactions between them that may be important for breast cancer prediction and possibly used as biomarkers. The Monte Carlo Feature Selection and Interdependency Discovery (MCFS-ID) algorithm was used to select top significant features that distinguish cancerous from normal tissue samples from TCGA data. The final feature set was derived by merging MCFS-ID outcomes from two independent steps (Fig 2). Firstly MCFS-ID was performed on the joined set consisting of mRNA and miRNA expression and DNA methylation. As a result, the feature ranking was dominated by the methylation features, followed by mRNA expression (Table 2, S1 Table). Only six miRNA expression features were found as relevant in breast cancer prediction. In the next step, three more MCFS-ID experiments were conducted on datasets consisting of single feature types to verify whether each of those is informative in distinguishing cancer from normal samples (S1 Table). With this approach, it was possible to expand the number of significant features, especially for miRNA data, and confirm the statistical significance of each individual set of attributes in sample classification. Finally, out of 417,486 multi-omics input features, 2,701 (2006+590+105) of them (Table 2) were selected by the algorithm as features potentially involved in cell physiological changes resulting in breast cancer development. The last two columns in Table 2 show significantly high weighted predictive accuracy (wAcc) of support vector machines (SVM) and random forest (RF) models, where the test samples were not used in the feature selection phase (Fig 1). It is worth underlining that after the MCFS-ID run, each feature is evaluated by the RI (relative importance) value so for each data type, instead of an unordered set of features, a ranking in which top features are the most informative is produced (S1 Table).

**Table 2.** Number of significant features returned by the MCFS-ID rankings together with RF and SVM results for classification of samples into cancer vs normal, shown as weighted accuracy (wAcc), performed using the features pointed in the “Sum”.

|  | Joined-set | Individual-set | Intersection | Sum | RF wAcc | SVM wAcc |
| --- | --- | --- | --- | --- | --- | --- |
| DNA methylation | 1504 | 1987 | 1485 | 2006 | 0.9068 | 0.9398 |
| mRNA | 432 | 588 | 430 | 590 | 0.9347 | 0.9347 |
| miRNA | 6 | 105 | 6 | 105 | 0.9848 | 0.9370 |

### Descriptive analysis of mRNAs having a significant predictive value

The Machine Learning feature selection process is focused on selection of features based on their high statistical significance, however it does not consider their biological meaning, which must be examined afterwards. At first, the top 10 mRNA genes from the MCFS-ID ranking (*ADAMTS5*, *COL10A1*, *TMEM220*, *ARHGAP20*, *MMP11*, *CAVIN2*, *PLPP3*, *MICU3*, *MME*, *CD300LG)* were screened and all of them were confirmed to have an association to cancer prediction and development. The top five were reported as effective cancerous tissue markers [60]. There is a number of scientific research, for each of the top 10 mRNA genes, well documenting their significance and association with cancerogenesis: *ADAMTS5* [61], *TMEM220* [62], *ARHGAP20* [63], *MICU3* [64]; or precisely with breast cancer: *COL10A1* [65], *MMP11* [66], *CAVIN2* (formerly known as *SDPR*) [67], *PLPP3* [68], *MME* [69], *CD300LG* [70], confirming the potential usefulness of the implemented approach.

Subsequent analysis included all significant mRNA genes to unveil their biological role in cancerogenesis and to discover new significant bio-functional relationships. Among 590 mRNA genes returned by MCFS-ID, 576 revealed differential expression, when filtering by the required log2FC and adjusted p-value (Fig 4A). Interestingly, these differentially expressed genes (DEGs) seem to be strong singularly independent predictors of breast cancer. At the level of feature selection performed with MCFS-ID, the returned decision trees (see Methods 2.2) presented very shallow depth and classical statistical tests confirmed that these genes significantly differed in expression between cancer/normal samples. Moreover, the majority of DEGs demonstrated a lowered expression in cancer (n=447), whereas only 129 showed increased expression. The down-expressed genes were enriched in 16 pathways from the Reactome database [71] that showed a great functional heterogeneity (Fig 4B), whereas over-expressed genes were enriched in 91 pathways (Fig 4C and S2 Table), in majority related to the cell cycle and mitosis. Down-expressed DEGs were enriched in pathways related to lipids regulation and transport, neurotransmission and retinoic acid synthesis (Fig 4B). These findings were confirmed by a Natural Language Processing approach (NLP). The pathways in which DEGs were enriched correspond very well to the unique keywords that describe clusters built on the gene function descriptions using NLP methods and hierarchical clustering (see “Detection of significant features using MCFS-ID algorithm”). The two most numerous clusters (mostly down-expressed) were related to the following terms: ‘*the regulation and metabolic processes’* and ‘*ion transport’* (Table 3). Finally, we confirmed that the set of 590 mRNA genes was significantly enriched in genes related (n=79) to immunological processes (chi-squared test, p<0.05). This fact confirms well known engagement of immune related genes in cancerogenesis.

**Fig 4.**
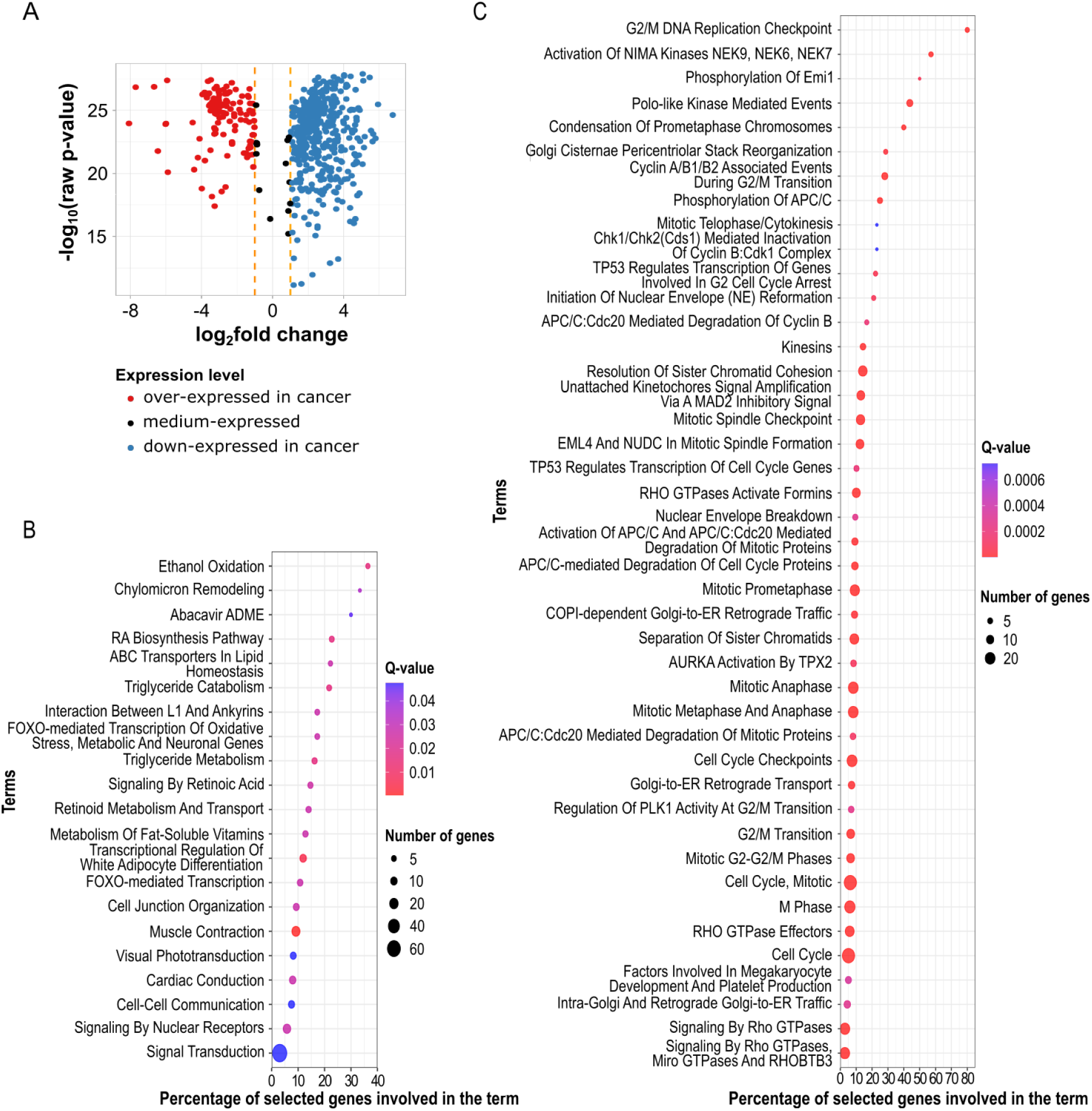
Overview of mRNAs indicated as significant in distinguishing cancer from normal tissue samples (A) The volcano plot shows differences in the expression levels of 590 mRNA considered significant in the cancer/normal prediction in the feature selection set (adjusted raw p-value>0.05). (B) Enriched pathways from the Reactome pathway database for down-regulated genes. (C) Enriched pathways from the Reactome pathway database for over-expressed genes. To allow for better readability, the number of less enriched pathways in the graph was reduced with the cutoff q-value=0.001 (all terms for cutoff q-value=0.05 are available in S2 Table).

**Table 3.** The result of NLP clustering of the significant mRNA genes: gene clusters, their most relevant keywords and gene expression statistics.

| Cluster ID | Cluster size | Top words associated with genes in cluster | Keywords interpretation | Mean log fold change for verification set | Direction of change in expression |
| --- | --- | --- | --- | --- | --- |
| 1 | 393 | regulation, process, metabolism, negative, negative_regulation, response, positive, positive regulation, gene, metabolic | regulation and metabolic processes | 1.5894 | over-expressed: 78<br>down-expressed: 313 |
| 2 | 38 | transport, ion, transmembrane, transmembrane_transport, calcium, abc, muscle, cardiac, ion_transmembrane, contraction | ion trans-membrane transport | 2.5636 | over-expressed: 4<br>down-expressed: 34 |
| 3 | 18 | receptor, g, coupled, g_protein, protein_coupled, gpcr, coupled_receptor, receptors, protein, ligand | receptor proteins | 2.8793 | over-expressed: 0<br>down-expressed: 18 |
| 4 | 24 | transcription, polymerase, rna_polymerase, rna, polymerase_ii, ii, regulation_transcription, transcription_rna, differentiation, development | transcription process | 1.5496 | over-expressed: 4<br>down-expressed: 20 |
| 5 | 31 | mitotic, cell_cycle, cycle, g, cell, apc, transition, apc_c, c, g_transition | cell cycle regulation | -2.5362 | over-expressed: 28<br>down-expressed: 3 |
| 6 | 16 | golgi, transport, er, golgi_er, retrograde, vesicle, vesicle_mediated, mediated_transport, mediated, traffic | golgi apparatus related | -0.9197 | over-expressed: 11<br>down-expressed: 5 |
| 7 | 4 | biological_process, biological, process | biological processes | 2.7369 | over-expressed: 0<br>down-expressed: 4 |

The Interdependence Discovery in MCFS-ID algorithm enables the additional identification of interdependencies between input features. These interdependencies are presented in a form of directed graph that visualizes knowledge derived from thousands of decision trees, highlighting feature RI with color intensity, interaction frequency with node size, and interaction strength and direction with edge/arrow thickness [7]. We focused on the ID-graph (S2 Fig) representing interdependencies between mRNA features due to their well-established functional descriptions. Among the strongest associations, we selected a few to demonstrate their biological relevance. For example, the *SRPX* → *HPSE2* interdependency suggests a link between the *SRPX*-encoded extracellular matrix protein, involved in cell adhesion, proliferation regulation, and potentially phagolysosome assembly and apoptosis [72], and the *HPSE2*-encoded heparanase enzyme, which degrades extracellular matrix proteoglycans and may contribute to angiogenesis and tumor progression [73]. Both genes potentially affect extracellular matrix-related biological processes. Another gene pair interdependency, *PLSCR4* → *PDGFA*, is known to play a functional role in apoptosis [74]. PDGFA connects with ALDH1L2 through PLSCR4, suggesting that *EMP2* overexpression activates *PDGFA* via the ALDH1L2-PLSCR4 pathway, initiating fibrin clot and apoptosis recognition [75]. Additionally, one may notice another interdependency of *PDGFA* with another feature *RNF186*, a component of an endoplasmic reticulum stress-activated apoptotic signaling pathway [76] and a well-known biomarker in several cancers, especially breast invasive carcinoma (BRCA) [77]. Here the functional association of the two features in the apoptosis seems reasonable. The final interdependency highlighted in our analysis, to be presented here, underscores the significance of *LYVE1* and *ADGRD2*, both membrane-associated features. Recent research by Anstee et al. [78] emphasized the role of LYVE-1+ macrophages in forming a CCR5-dependent perivascular niche linked to immune exclusion and therapy resistance in murine breast cancer. Whereas, the *ADGRD2*, a G protein-coupled receptor, is known to contribute to various aspects of cancer, including proliferation, angiogenesis, invasion, and metastasis [79]. This short analysis of interdependencies shows that ID graphs (S2 Fig) seem to be a rich source of new useful information, however contextual influence and interactions between various biological mechanisms/factors is a part of another extremely complex analytical layer that is not in the main focus of this paper.

### Genomic context of DNA methylations with predictive value

Feature selection revealed 2006 significant sites that differed in methylation levels between cancer and normal samples (Table 2), hereafter called differentially methylated sites (DMSs). Out of all DMSs only one locus (cg02025583) is located within the promoter of one of the top 10 mRNA genes returned in the main MCFS-ID ranking. This promoter precedes *TMEM220* gene and the cytosine cg02025583 is overlapped by a motif of the E2F2 transcription factor (TF), which is a good example of altered epigenetic regulation of gene expression.

The distribution of Illumina Infinium Human Methylation 450K BeadChip probes in genomic regions (Illumina 450K array) was compared with the distribution of DMSs and DMSs were found statistically significantly enriched in CpG Islands (CpGI) and open seas as well as their depletion in shores (chi-squared test, corrected p-value≤0.05, Fig 5A). The indicated sites are candidates for modulating activity of regulatory regions therefore we focused on their methylation levels in normal vs cancer samples to unveil their putative regulatory role in cancer development. We found that DMSs methylation *β*-values showed a significant shift towards higher values within CpGI and presented significantly different distribution of *β*-values within shores and open seas between cancer vs normal (Wilcoxon test p-value≤0.05, Fig 5B). There were over two times more hyper-methylated DMSs (n=479) than hypo-methylated DMSs (n=225) discovered in tumors (Fig 5C). Interestingly, the vast majority of hyper-methylated DMSs were located within CpGI, which are well known gene transcription regulators (Fig 5D). Medium methylated DMSs showed very high frequency not only in CpGI but also in shores, shelves and open seas (Fig 5D).

**Fig 5.**
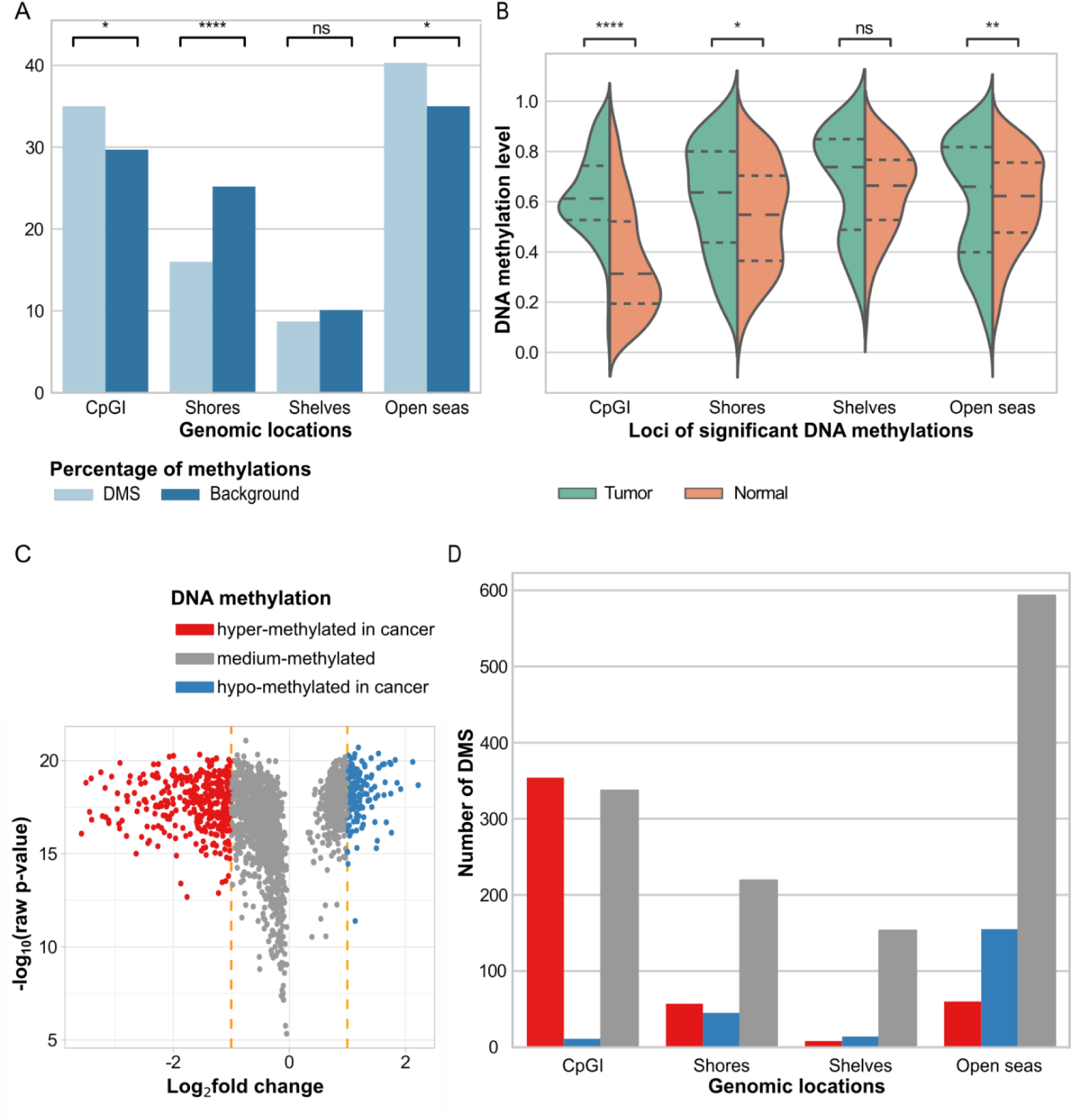
Characteristics of 2006 significant DNA methylation sites (DMSs). (A) Distribution of loci included in the Illumina 450K array in comparison to the distribution of DMSs. (B) DMSs *β*-values distribution in the tumor and normal samples with respect to the specific genomic regions. (C) DMSs assigned to hyper/medium/hypo-methylated with respect to log2 Fold Change of *β*-values. (D) Number of loci with hyper/medium/hypo-methylated DNA in particular types of genomic regions.

To investigate a potential regulatory association of gene expression mediated by DNA methylation, the Spearman correlation was measured between mRNA levels of 590 significant genes and *β*-values of each DMS within 1 Mbp upstream and downstream from TSS of these genes. The correlation cut-off value was set to |*rho*|≥0.6 and there were 59 pairs meeting this condition (S3 Table). The majority of the obtained correlations were negative (n=44) with only a few positive (n=15). More frequent negative correlation was expected, if we assume that DNA methylation located in promoters or enhancers inhibits gene expression. Among these pairs there were 34 unique genes and 55 unique DMS loci. Almost all genes were down-regulated in tumor samples, but the *HN1L* and *KIFC1* were up-regulated. Out of 55 DMSs five were located within one gene promoter. Interestingly, all of them were hyper-methylated and were close to each other, within one CpGI in a range of 25 bp, within the *NKAPL* gene promoter (S3 Fig). Two of them (cg18694169, cg10253847) were overlapped by a motif of the NRF1 TF. NRF1 normally activates gene expression, but here, due to hyper-methylation, its binding to DNA would be inhibited. Accordingly, *NKAPL* was down-regulated in cancer samples, which could be explained by hyper-methylation of the five DMSs [80]. The other 50 DMSs were not assigned to any promoter and also had a confirmed chromatin state indicating possible activity (S3 Table), so they were defined as potential distal regulatory factors.

In order to have a better understanding of a potential role of the DMSs in gene expression regulation, all DMSs were intersected with chromatin states of MCF-7 breast cancer cell line (Fig 6A) and simultaneously all sites from Illumina 450K were intersected with chromatin states of MCF-7 as well (Fig 6B). The observed distribution of DMSs across chromatin states was similar to the distribution of all Illumina 450K sites across the chromatin states (chi-squared test p-value>0.2). Comparison of the distribution of hypo- and hyper-methylated DMSs revealed a visible enrichment of hyper-methylated DMSs in transcriptionally inactive chromatin states, like heterochromatin. At the same time, hyper-methylated DMSs were depleted within states associated with gene transcription activation, such as enhancer, promoter and transcribed regions (Fig 6C). Conversely, for the hypo-methylated DMSs the opposite pattern was observed (Fig 6C).

**Fig 6.**
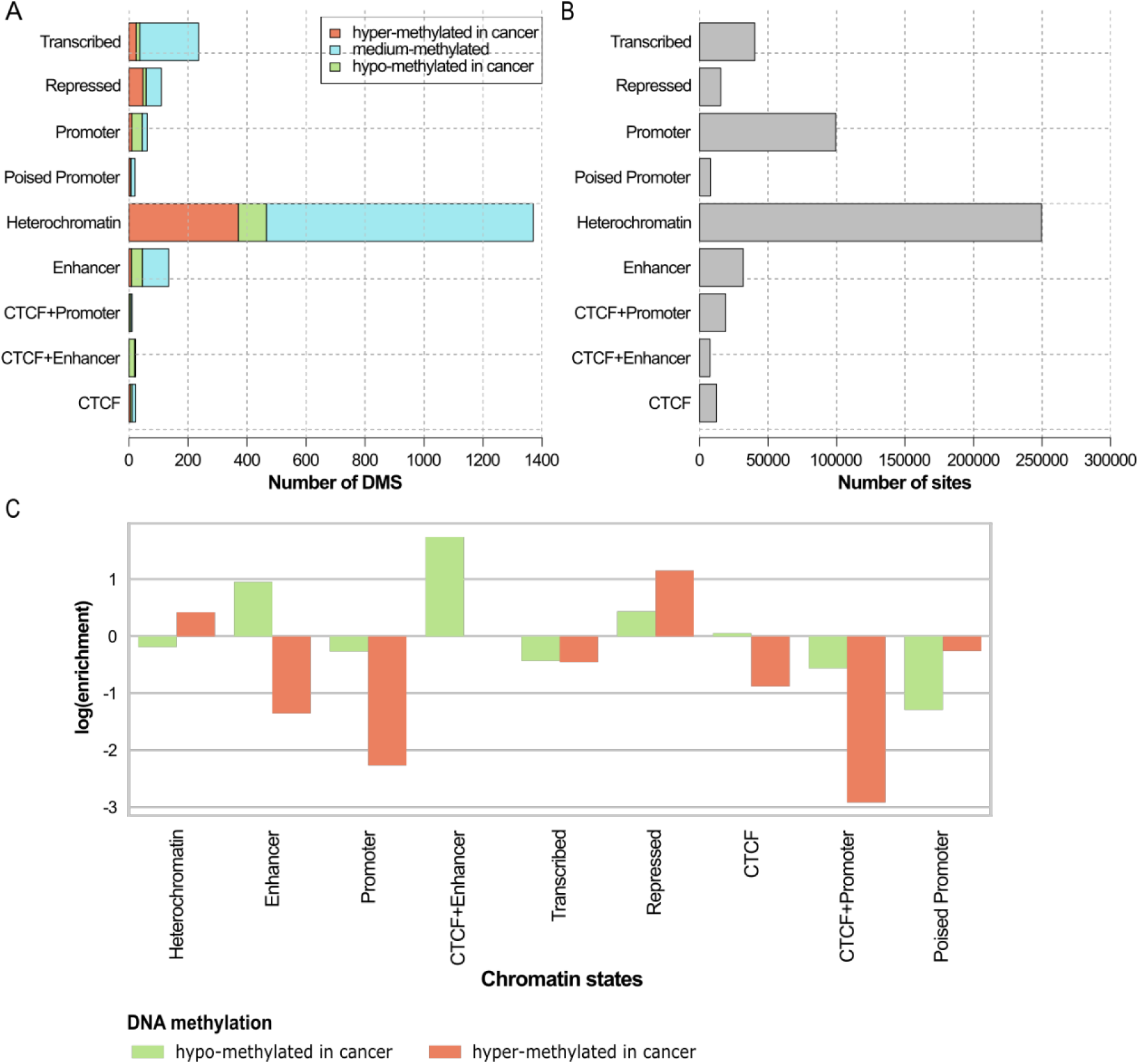
Distribution of cytosines across chromatin states obtained for the MCF-7 breast cancer cell line. (A) Number of DMS in individual chromatin states. (B) Illumina 450K sites assigned to individual chromatin states, representing the background distribution. (C) Differential distribution of hypo- and hyper-methylated DMS in specific chromatin states.

Next, the evaluation of the impact of DMSs on survival was tested. Out of 2006 DMSs there were 691 (S4 Table) that were found to be significantly correlated with patients survival (p<0.05). The number of loci associated with survival in 100 random picks of 2006 methylation probes was between 375 and 450, which confirms that MCFS-ID returned an enriched list (S4 Fig). However, after application of the correction for multiple testing, none of the sites achieved expected statistical significance. Therefore, the prediction of patients’ survival by the DMSs should be treated with caution.

### Biological role of significant miRNA genes

To characterize the regulatory functions of 105 miRNAs identified in the MCFS-ID experiment, their associations with mRNA from the miR+Pathway database were verified, resulting in detection of 822 unique mRNAs. Out of these mRNAs, 43 were shown to be significant in predicting breast cancer, in the MCFS-ID analysis (results section “Detection of potential breast cancer biomarkers using the MCFS-ID algorithm”). The intersection with the mRNA clusters returned by NLP analysis showed that out of 43 mRNAs, 32 belong to cluster 1, one mRNA to cluster 2, two to cluster 3, seven to cluster 5, and one to cluster 6 (Table 3). Most of the mRNAs assigned to cluster 1 had decreased expression in the cancer samples but five had increased expression. The opposite situation can be observed in cluster 5 where six mRNAs were upregulated in the tumor and one down-regulated.

For 58 miRNAs genes over-expressed in cancer, selected out of the 105 significant miRNAs (S1B Fig), the target mRNA genes were assigned to them and the putative associations between 46 miRNAs and 126 mRNAs were confirmed with Spearman correlation (*rho* ≤ -0.2, S5 Table). KEGG pathway analysis of these 126 mRNAs returned insignificant results (adj. p-value > 0.05). The mRNAs contributing to significant correlations formed a protein-protein interaction network consisting of 2265 proteins. For the 50 proteins with the highest number of interactions in this network, the miRNAs targeting their mRNAs were assigned (S6 Table), resulting in 22 unique miRNAs, all initially obtained as significant in the main MCFS-ID run. KEGG pathway analysis of the genes encoding those 50 proteins showed significant annotations to breast cancer (p-value = 0.000454), as well as to many other cancers e.g. melanoma, renal cell carcinoma, acute myeloid leukemia, colorectal cancer (S5A Fig). These proteins were also found to be associated with cancerogenesis-related biological processes, e.g. chemical, viral or proteoglycans, as well as pathways known to be crucial for cancer development, e.g. p53 signaling pathway (S5A Fig). The returned terms from GO BP analysis were almost all related directly or indirectly to cell cycle, the crucial process for cancer development and progression (S5B Fig). Moreover, out of the 50 proteins with the highest number of interactions in the miRNA-regulated protein-protein network, 16 are known drug targets in breast cancer treatment. The largest number of them was targeted by palbociclib, ribociclib [81] and abemaciclib [82], among 19 others drugs (S7 Table). It is worth mentioning that both scores showed in S7 Table, i.e. Drug Score (DScore), which measures the suitability of the drug according to the genomic profile, and Gene Score (GScore), which reflects the biological relevance of genes in the tumoral process, had high values for the majority of aforementioned drugs, indicating significant effect of these drugs (S7 Table). The resulting miRNA-Protein-Drug network is visualized in the S1C Fig There are two mRNA genes *CDC25A* and *BIRC5* specified in this network, whose expression significantly correlated with upregulated mi-RNAs, namely hsa-mir-100, and hsa-mir-218-2, respectively, that are also known to be breast cancer drug targets (S7 Table, S5A,B Fig) These two miRNAs were selected by MCFS-ID main run.

### Detection of miRNA and DNA methylation loci significant in the context of predicting mRNA expression levels

The result of 590 MCFS-ID experiments run on 590 significant mRNA features (Table 2), each separately used as the target variable, with miRNA expression or DNA methylation set as predictors showed that miRNA features are better predictors as compared to methylation. Out of 590 mRNA expression features, only 73 can be correctly predicted by miRNA features, 66 by DNA methylation features, and 39 by both (where Pearson correlation level ≥ 0.8). For each target variable (out of 590) a different significant set of features was returned. These sets differed in size and contained different features in the top ranking, therefore it was possible to analyze the distribution of: significant set size (based on MCFS-ID cutoff), Pearson correlation calculated between each mRNA expression and its prediction based on the significant feature set. To find out if, for different mRNA, there are common predictive miRNAs or DNA methylations the frequency of a single significant feature across all significant feature sets was calculated as well (separate for miRNA and DNA methylation). Histograms show that the number of selected significant DNA methylations in the rankings was much greater than for miRNA (see Fig 7A,B), which may correspond to the size of the input data sets and to the fact that statistical modeling of mRNA expression is much more complex in the case of DNA methylation data. However, the quality of the prediction of mRNA expression, based on miRNA features (see Fig 7C,D) is comparable to that achieved with the help of DNA methylation data.

**Fig 7.**
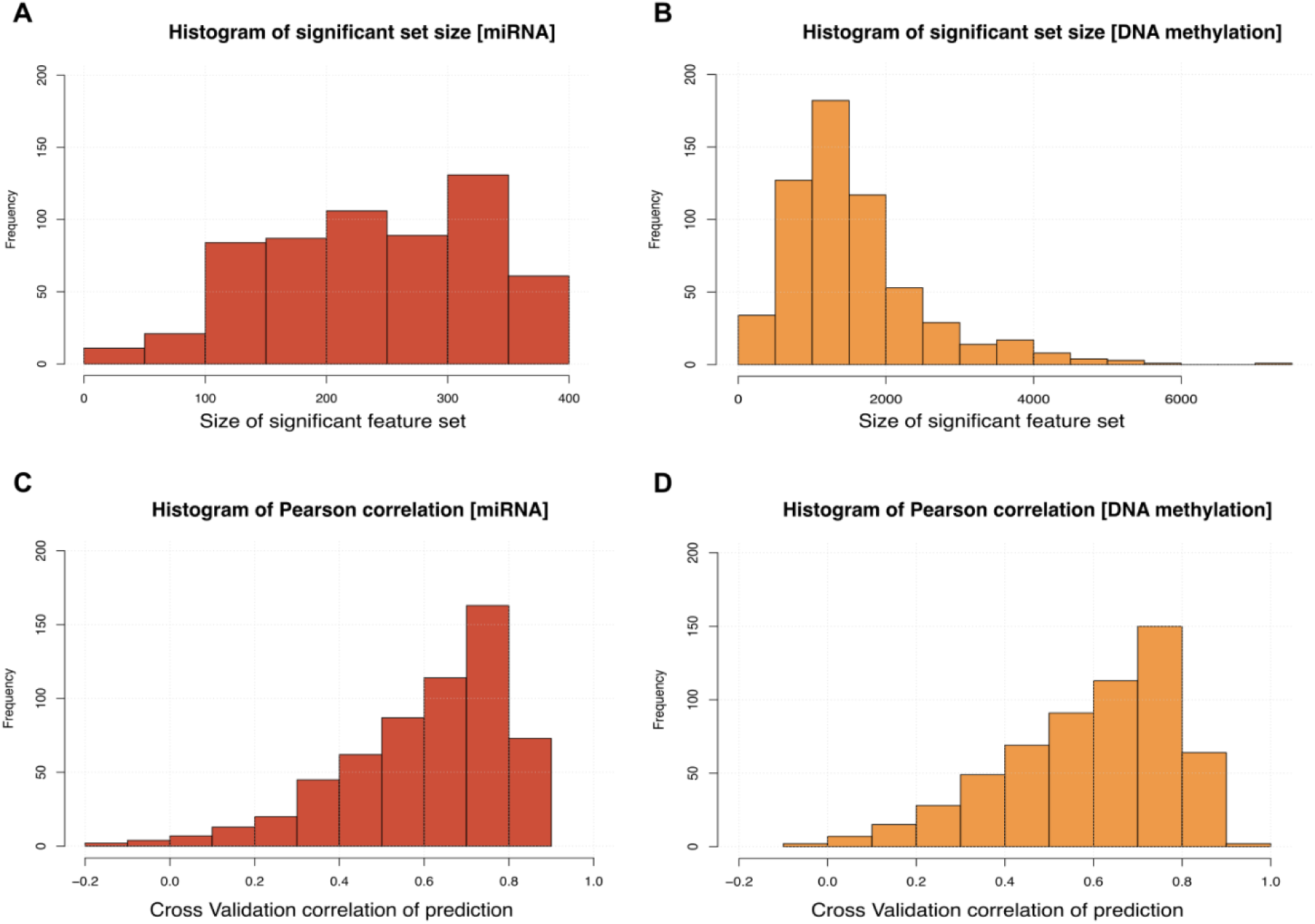
Summary of mass MCFS-ID experiments on 590 mRNA genes. (A) Distribution of the number of significant miRNA features returned across 590 experiments. (B) Distribution of the number of significant DNA methylation loci returned across 590 experiments. (C) Distribution of the Pearson correlations obtained for linear models built on significant miRNA features returned across 590 experiments (D) Distribution of the Pearson correlations obtained for linear models built on significant DNA methylations returned across 590 experiments.

To get a better overview of all the selected top features, all MCFS-ID top rankings (separated by data category) were combined and, for each predictor feature, a sum of RI, mean RI, and frequency (how many times a single feature was found as significant) were calculated. The resulting two rankings - separate for miRNA expression and DNA methylation are shown in Table 4 (the top 15 features) and supplementary material (S8 Table, S9 Table). For 73 mRNA target features (that could be predicted with Pearson correlation level ≥ 0.8), 97 miRNA predictors were found by MCFS-ID as significant for all 73 mRNAs features, which means that the same miRNAs take a part in a successful prediction of expression of the 73 protein coding genes. The predictive impact of particular miRNAs changes depending on mRNA, which is observed as a different position in a single MCFS-ID ranking. Moreover, 81 out of 97 miRNA genes were also significant in cancer prediction and selected by the main MCFS-ID experiment (see section “Detection of potential breast cancer biomarkers using the MCFS-ID algorithm” and Table 2). The last column in Table 4 refers to the ranking of the main MCFS-ID experiment (on a given data type) and all the top 15 miRNA genes in the table are confirmed as cancer specific in literature according to www.mirbase.org.

**Table 4.** Top 15 miRNA genes (left table) and DNA methylation (right table) loci from the mass MCFS-ID experiments where mRNA expression was set as a target variable and prediction of the regression tree on the top significant set provided Pearson correlation ≥0.8.

| miRNA gene | Freq | Sum RI | Mean RI | MCFS-ID rank | DNA Methylation | Freq | Sum RI | Mean RI | MCFS-ID rank |
| --- | --- | --- | --- | --- | --- | --- | --- | --- | --- |
| hsa_mir_139 | 73 | 65.628 | 0.899 | 1 | cg07267550 | 7 | 3.044 | 0.435 | 1234 |
| hsa_mir_141 | 73 | 38.519 | 0.528 | 10 | cg00914963 | 7 | 3.036 | 0.434 | 2048 |
| hsa_mir_10b | 73 | 36.690 | 0.503 | 2 | cg19533977 | 7 | 3.024 | 0.432 | 25 |
| hsa_mir_183 | 73 | 33.020 | 0.452 | 4 | cg08113562 | 7 | 2.714 | 0.388 | 12061 |
| hsa_mir_140 | 73 | 32.825 | 0.450 | 11 | cg17901038 | 7 | 2.620 | 0.374 | 186 |
| hsa_mir_200a | 73 | 31.182 | 0.427 | 15 | cg18253799 | 7 | 2.584 | 0.369 | 2217 |
| hsa_mir_96 | 73 | 28.417 | 0.389 | 8 | cg20417953 | 7 | 2.504 | 0.358 | 6490 |
| hsa_mir_429 | 73 | 25.452 | 0.349 | 20 | cg20524128 | 7 | 2.488 | 0.355 | 2066 |
| hsa_mir_204 | 73 | 23.546 | 0.323 | 12 | cg10520594 | 7 | 2.393 | 0.342 | 4407 |
| hsa_mir_99a | 73 | 23.041 | 0.316 | 6 | cg20701457 | 7 | 2.159 | 0.308 | 1487 |
| hsa_mir_592 | 73 | 22.352 | 0.306 | 16 | cg16009970 | 7 | 2.090 | 0.299 | 13163 |
| hsa_mir_378 | 73 | 22.320 | 0.306 | 38 | cg06976025 | 7 | 2.012 | 0.287 | 3878 |
| hsa_mir_145 | 73 | 22.218 | 0.304 | 5 | cg15601264 | 7 | 1.993 | 0.285 | 188 |
| hsa_mir_21 | 73 | 21.700 | 0.297 | 3 | cg22608492 | 7 | 1.932 | 0.276 | 151 |
| hsa_let_7c | 73 | 21.686 | 0.297 | 13 | cg11441693 | 7 | 1.913 | 0.273 | 12369 |

### Tracking associations between DMSs and detected TF motifs

Depending on the cytosine location in the genome or changes in its DNA methylation level, the cytosine loci overlapping transcription factor binding sites (TFBS) may significantly affect binding affinity of a TF to the DNA. To investigate this issue, for each DMS a DNA sequence covering 41 bp was obtained (site +/- 20 bp) and TF motif search was applied. First, DMSs were divided according to their location in genomic regions: promoters, gene bodies and intergenic. There were 616 DMS located within promoters, 1037 in gene bodies and 353 in intergenic regions, and the motif search returned 48, 54, 21 TF motifs in these three genomic regions, respectively. The numbers of returned TF motifs proportional to DMSs reflect no significant enrichment in the mentioned specific genomic regions (chi-squared test, p-value = 0.138). Moreover, it was confirmed that none of the protein families is overrepresented among motifs confirmed within the three genomic regions (p-value after FDR correction > 0.05). There were 45 common motifs between promoter and gene body regions of which 21 were also shared with intergenic. Specifically, we detected three TFs: E2F1, ESR2, NRF1 only in promoters and nine: ELK3, HIF1A, ITF2, LYL1, NFIA, NR2C2, SNAI1, ZN341, ZN589 only within gene bodies (S10 Table).

When verifying TF motifs overlapping hyper-methylated (n=479) cytosines, we confirmed 48 motifs and for hypo-methylated loci (n=225) there were 5 motifs detected. Only one motif, EPAS1_HUMAN.H11MO.0.B was specific for the hypo-methylated set of DMS and the remaining four were detected for both cytosine methylation levels (Fig 8A, B).

**Fig 8.**
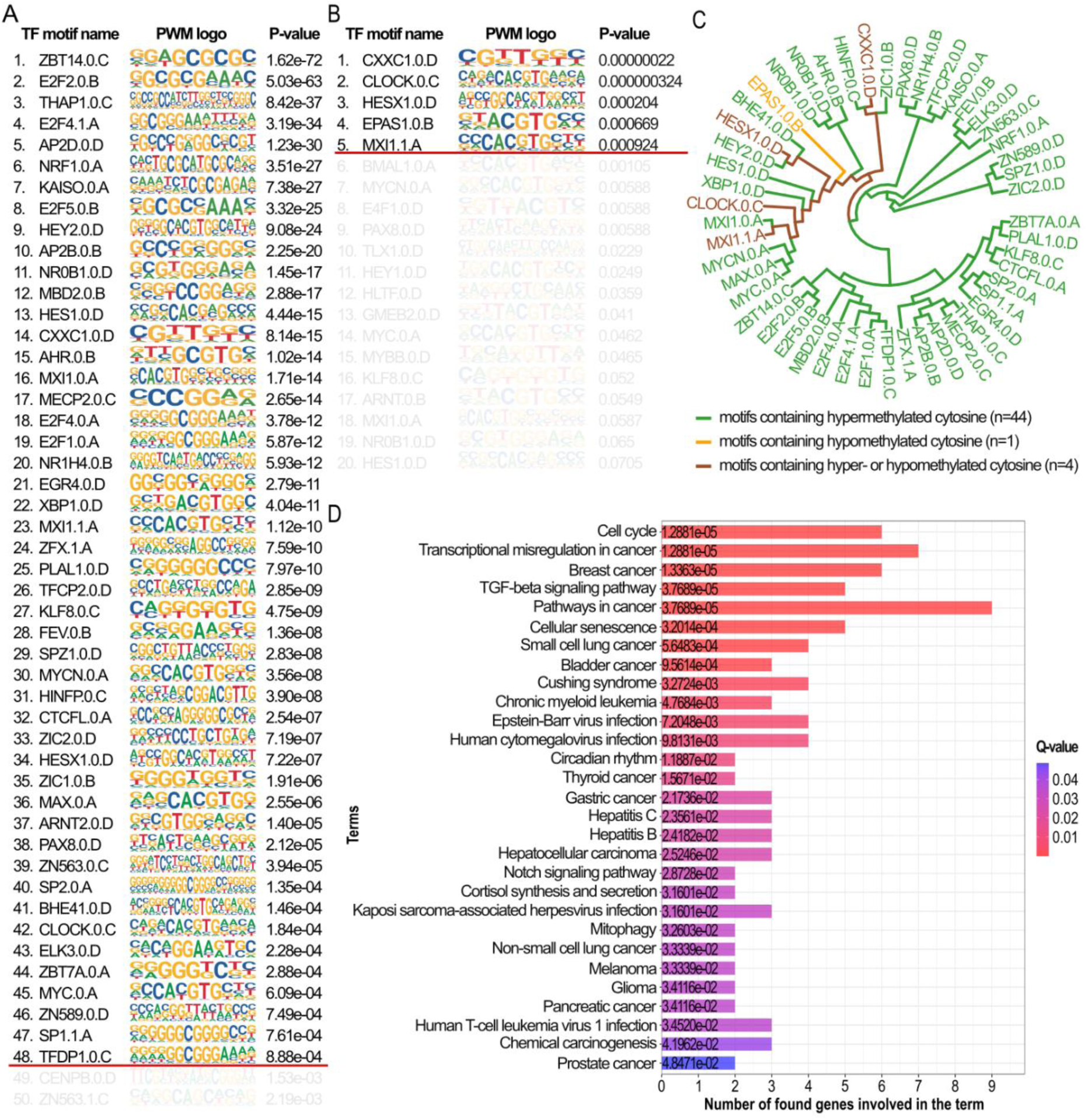
TF motifs overlapping differentially methylated cytosines. (A) TF motifs overlapping hyper-methylated DMS. (B) TF motifs overlapping hypo-methylated DMS. In (A) and (B), the red horizontal line indicates the p-value cut-off point. (C) Hierarchical clustering of TF motifs based on their PWMs. (D) Functional analysis of genes encoding TFs whose motifs overlapped DMSs hyper-methylated in cancer (KEGG database). The list of genes related with specific terms is shown in S11 Table.

To verify the similarities of PWMs of the detected TF motifs (Fig 8A, B) the hierarchical clustering was performed and it showed that the majority of motifs overlapping hyper-methylated cytosines (Fig 8A) constituted two homogenous clusters. The third cluster contained motifs overlapping both hypo- and hyper-methylated cytosines, therefore it seems that TF motifs have some common characteristics independent from cytosine methylation level. Next, using the KEGG database [83] (Fig 8D) it was verified that the majority of biological pathways related to genes encoding TF motifs overlapping hyper-methylated DMS were precisely connected with cancer (e.g. misregulation in cancer, breast cancer, lung cancer etc.) as well as through pathways which are well-known to be connected with tumorigenesis (e.g. TGF-β signaling pathway [84], human cytomegalovirus infection [85]). Of note is the group of genes *SP1*, *MYC*, *HEY2*, *E2F1*, *E2F2* and *HES1* known to be associated with KEGG breast cancer functional pathway (Fig 8D).

### Models of regulatory networks

To discover interdependencies between DMSs and genes encoding TF motifs overlapping these DMSs in the context of breast cancer prediction, another MCFS-ID experiment was conducted (S12 Table). Only the genes encoding TFs that were overlapping significant DMSs, as well as the levels of DNA methylations of these DMSs, were considered. The MCFS-ID algorithm returned 281 significant features (S13 Table), out of which the vast majority were DMSs with only 7 genes encoding TFs, namely: *MXI1, EPAS1, PLAGL1, E2F1, NR0B1, BHLHE41* and *ARNT2*. These 281 features were annotated to their closest genes resulting in identification of 279 mRNA genes named hereafter target genes. Nine of them: *PITX1*, *TGFBR3*, *PAFAH1B3*, *TAL1*, *TDRD10*, *SHE*, *LEP*, *TMEM220* and *NKAPL*, were previously reported in the main MCFS-ID ranking (S1 Table). Only one target gene: *PITX1,* contained a motif of one of the 7 TFs encoded by the significant genes identified in the second MCFS-ID experiment, namely *MXI1* overlapping hyper-methylated DMS cg00396667 cytosine and the remaining 8 target genes were linked only to the DMS, not to TFs. For breast cancer, it was confirmed that down-regulation of *PITX1* improves prognosis and it is associated with DNA methylation levels [69, 86].

To review possible regulatory dependencies, mRNA expression values of these 9 target genes were used as prediction variables in a set of linear models. As the result, four linear models for the following target genes: *TMEM220*, *NKAPL TGFBR3*, *SHE* reached statistical significance (p≤0.05, R^2^>0.5) and these genes were located on 3rd, 44th, 245th and 311th position in the main MCFS-ID ranking, respectively (S1 Table). All four linear models are highly impacted by tissue type value, however after removing this feature from the set of explanatory variables, prediction of the models holds on a relatively high level (Pearson correlation calculated between target gene expression and predicted value decreases from 0.7-0.8 to 0.6-0.7 after removing tissue type - see S14 Table). This observation suggests that the relation between target gene expression and linked TFs features is noticeably strong and specific in the context of tissue type. The four target genes are down-regulated in tumor samples, suggesting that they are tumor suppressors regulated by hyper-methylated DMSs that reduce TFs binding affinity. Moreover, these DMSs were in heterochromatin regions, which is in the line with gene silencing. To illustrate the hypothesis that DMSs located within the TF motif causes disruption of TF binding, we visualized features of linear models in a way that the symbols of genes that encode TFs were connected with DMSs and then with their target gene (Fig 9A). Additionally, based on the Spearman correlation results between the up-regulated miRNAs and their top target mRNAs (S5 Table), one additional association of hsa-miR-211 with *SHE* was detected and added to the visualization (Fig 9A).

**Fig 9.**
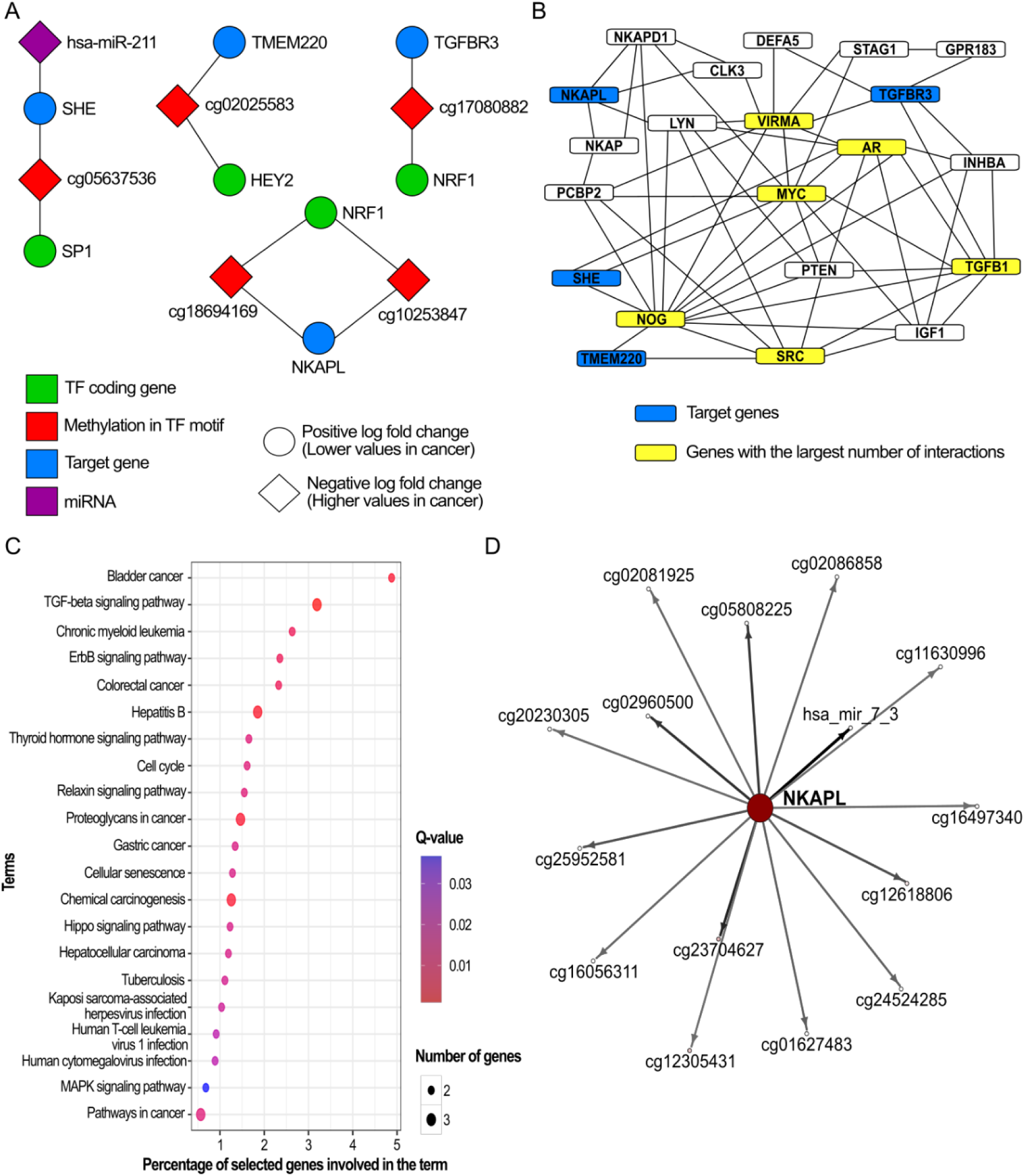
Target genes interactions, biological functions and graphical representation of their putative regulatory elements. (A) Visualization of interactions driven from linear models. (B) The network of gene-gene interactions created for the identified target genes visualized on A. (C) KEGG pathway analysis for 10 genes highlighted in B. (D) ID-graph for *NKAPL* gene.

Additionally, to obtain a better insight into tumor suppressive paths involving target genes, the gene-gene interactions were established using Pathway Commons and aggregated into one network (Fig 9B). At first, the direct interactions with a group of new genes were established and then the most frequent of these new genes were used as an input to Pathway Commons to obtain levels of indirect interactions. This approach helped to build a more dense network to select genes with the largest number of putative interactions: *MYC*, *NOG, TGFB1*, *VIRMA, SRC* and *AR* genes, pointing at their significance in this network. Genes: *NOG, TGFB1*, *VIRMA, SRC* were over-expressed in cancer, unexpectedly *MYC* was down-expressed and AR showed no significant difference in Wilcoxon statistical test (S6A Fig). Genes *MYC* and *AR* are known to be over-expressed in specific breast cancer subtypes [87–89], but this pattern was not proven for the entire, much more heterogeneous, cohort of TCGA breast cancer samples.

Biological pathway analysis of the aforementioned six genes, and target genes used in linear models (*TMEM220*, *NKAPL TGFBR3*, *SHE*) showed enrichment of biological pathways related to various cancer types (e.g. bladder cancer, chronic myeloid leukemia, proteoglycans in cancer) and to signaling pathways whose alterations are often associated with carcinogenesis e.g. TGF-β signaling pathway [84], erbB signaling [90] or relaxin signaling pathway [91] (Fig 9C).

Finally, the Interdependency Discovery function of the MCFS-ID algorithm was used to find statistically significant, nonlinear interactions between features that amplify each other in the classification task. S2 Fig in the supplementary materials provides four ID-graphs created for top 50 features and top 50 strongest links between the features. These figures were created as an additional result of four main MCFS-ID experiments (described in Fig 2). Fig 9D shows a part of the ID-graph created for the *NKAPL* gene and its neighbors, connected by edges that represent the power of interaction between them. Recently, this gene has been shown as a significant driver of cancer development [92], a prognostic marker [93] and an important factor associated with resistance to pharmacotherapy [94]. The visible directions of interactions show that the *NKAPL* gene expression plays the major role in classification of normal and tumor tissues and the remaining DNA methylation and miRNA features boost its predictive power. Notice that hsa_mir-7_3 is known as a significant factor contributing to cancerogenesis [95]. S6B Fig provides additional ID-graphs created for genes that were used as target genes in linear models: *TMEM220*, *TGFBR3*, and *SHE* genes.

### Epigenomic Regulatory Spatial Model

The expression of a gene is directly associated with the distances between its body and regulatory regions, and these distances differ in three-dimensional space as compared to linear space [96]. Therefore, measurable and quantitative variations in spatial distance are subsequently responsible for changes in gene expression. Epigenetic processes, such as DNA methylation, impact the transcription machinery to influence gene expression [97]. Therefore, the spatial proximity of genes to some of the features identified as significant in the main MCFS-ID analysis, as discerning between breast cancer and normal samples, was verified at the level of 3D chromatin structure.

In the mass MCFS-ID analysis, several genes, whose expression was well predicted by DNA methylation levels, were found. One of them was *FXYD1* which was selected as an example to visualize in 3D the putative functional association between regulatory features and mRNA expression. (Fig 10A.i). To achieve this, an ensemble of 100 models was built using the 3D-GNOME approach [98, 99]. From this ensemble, the most representative spatial conformation was selected for visualization purposes in UCSF Chimera [100]. The gene promoter region and cg23866403 DNA methylation loci were observed to be in close proximity to the enhancer of the *FXYD1* gene in the normal sample, as compared to the longer distance between the regulatory elements in breast cancer, where *FXYD1* gene is down-regulated and cg23866403 loci is hyper-methylated, as shown in the box plots (Fig 10A.ii). The spatial distance distribution between enhancer-promoter and methylation site-promoter was also calculated, to see how these distances vary within the ensemble of 100 models (Fig 10A.iii).

**Fig 10.**
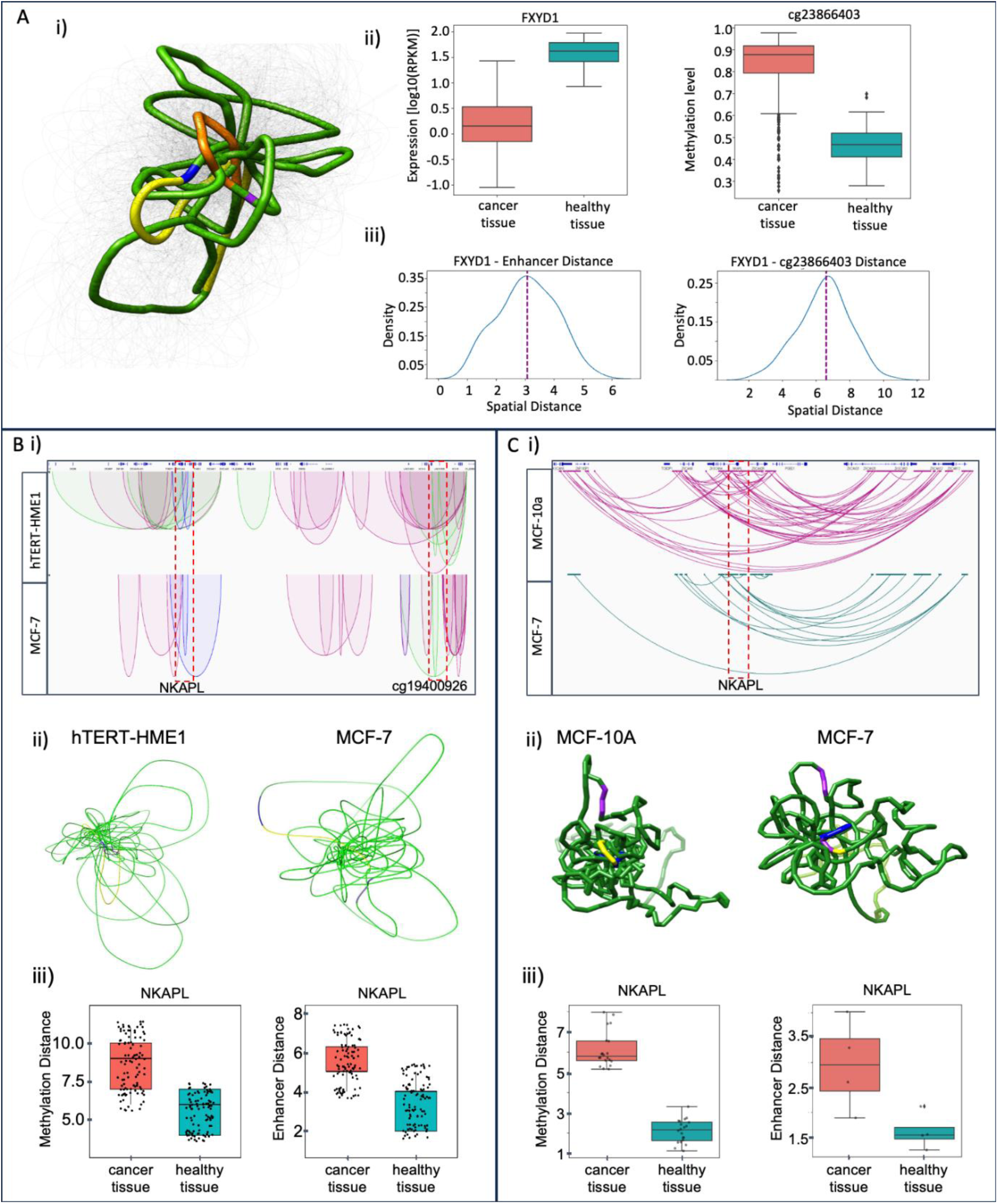
Spatial Regulatory Model of chromatin – (A) (i) The most representative chromatin 3D computational model from the ensemble of 100 spatial models generated by the 3D-GNOME method for the *FXYD1* gene with labeled promoter (blue), gene body (yellow) cg23866403 methylation loci (purple) and potential enhancer region (orange). (ii) The box plots show cancer and healthy samples *FXYD1* expression (left) and cg23866403 loci methylation levels (right). (iii) The spatial distance distribution between the *FXYD1* gene promoter and its enhancer region (left) and the cg23866403 methylation loci (right). (B) (i) Cohesin-mediated chromatin interactions around the *NKAPL* gene in the integrative genomics viewer for hTERT-HME1 (healthy) and MCF-7 (cancer) cell lines. Green color annotates enhancer-promoter loops, blue color promoter-promoter loops. (ii) The representative chromatin 3D model based on Cohesin ChIA-PET data for the *NKAPL* gene. (iii) The spatial distances between promoter-methylation (left) and promoter-enhancer (right) for cancer and healthy cell lines. (C) (i) PCHi-C interactions around the *NKAPL* gene in the integrative genomics viewer for MCF-10A (healthy) and MCF-7 (cancer) samples. (ii) Chromatin 3D model of the *NKAPL* gene in MCF-10A (left) and MCF-7 (right) cell lines. (iii) The spatial Euclidean distances between the *NKAPL* gene body and DMS (left); the *NKAPL* gene body and the enhancer (right) both for MCF-7 and MCF-10A.

To thoroughly investigate the contrast between the 3D structure of cancer and normal cells, a model was constructed around the *NKAPL* gene, which contributes considerably to the classification of normal and tumor tissues [80, 93] (also shown in the previous section “Models of regulatory networks”). To do this, two specific datasets - ChIA-PET and PCHi-C were considered. ChIA-PET data identifies 3D contacts (in this particular experiment mediated by the RAD21 protein), which provides a chromosome-wide 3D view of the target gene and its spatial connectivity. PCHi-C experiments provided a promoter-centric view of a target gene, with interactions between each gene promoter regions and other distal DNA segments, including regulatory elements.

Cohesin-mediated chromatin loops were explored by application of chromatin interaction analysis with use of paired-end tag sequencing data (ChIA-PET) downloaded from Encode [58] for two cell lines: MCF-7 (cancer) and hTERT-HME1 (normal); aligned to hg38 reference genome. It was confirmed that the identified cohesin-mediated loops (significant interactions) connecting two distant genomic fragments (anchors) surrounding the *NKAPL* gene were stronger and higher in number in hTERT-HME1 (normal) as compared to the MCF-7 (cancer). Moreover, visual examination of the loops in the genome browser illustrates that loops anchored by distal enhancers were confirmed only for hTERT-HME1 (Fig 10B.i). These enhancer-specific interactions in the hTERT-HME1 cell line work together with the promoter to control the expression of the *NKAPL* gene, and they are absent in the MCF-7 cell line (Fig 10B.i). Additionally, 3D models from ChIA-PET were constructed using 3D-GNOME algorithm to examine and annotate the resulting structures (Fig 10B.ii). We generated a spatial model of the NKAPL gene region for both cell lines by 3D-GNOME, and these models suggest that chromatin in this regions is more condensed in the normal cell line compared to the cancer cell line, in which it is loosely packed in three-dimensional space (Fig 10B.ii).

Next, the spatial distances in Euclidean space between: (1) the *NKAPL* gene body and DNA methylation sites or (2) *NKAPL* gene body and enhancer were calculated for the two cell lines MCF-7 and hTERT-HME1 (Fig 10B iii). These distances were obtained by mapping the beads from the 3D model representing gene body (*NKAPL*), enhancer (*NKAPL*) and DMS (S15 Table, S7 Fig) and calculating them for respective pairs of beads. The hypothesis that chromatin around this gene is more loosely packed in the cancer cell line than in the non-cancer one was confirmed by the box plots of the distance distributions (Fig 10B.iii, S7 Fig) for MCF-7 and hTERT-HME1 Cohesin ChIA-PET interaction sets.

To further investigate the differences between the 3D structure of this locus in cancer and normal cells, we examined promoter interactions with distal regulatory elements using PCHi-C datasets obtained from Javierre et al. [101] and Beesley et al. [59,102]. In PCHi-C, 23 interactions around the *NKAPL* gene in MCF-7 and 91 interactions in MCF-10A (normal tissue) were detected (Fig 10C.i). In this case 3D models were constructed using the Spring Model, which is based on the OpenMM molecular dynamics simulation engine based on a beads-on-string representation [57] (Fig 10C.ii). Additionally both 3D models are showing the loci with differential DNA methylation around the *NKAPL* gene reflecting higher DNA methylation in cancer (S15 Table, S7 Fig). Again, the distributions of the Euclidean distance (calculated as previously for ChIA-PET data) confirmed the hypothesis that chromatin in this locus is more loosely packed in cancer than in normal cell lines (Fig 10C.iii). The high concordance between the resulting box plot for PCHi-C and that of the ChIA-PET dataset confirms the reliability of the previous result.

## Discussion

In this study, the MCFS-ID algorithm was applied to return a ranking of statistically significant molecular features distinguishing cancerous and normal tissue samples deposited in The Cancer Genome Atlas (TCGA) [https://www.cancer.gov/tcga]. Using this algorithm we could select a small number of multi-omics features significantly different between cancer and normal, reducing the dimensionality of these datasets from 417k initial features to only 2.7k of ranked features. Our further effort focused on verifying whether these significant features also have a substantive meaning in the context of cancerogenesis. It was shown that almost all (n=590) mRNA significant genes returned by MCFS-ID reflected differential expression (Fig 4A) between cancer and normal samples. Nevertheless MCFS-ID also explores the interactions, therefore, the statistical significance of individual features for the entire group of samples is not obvious. Meanwhile, a large part of the DMSs did not show differential methylation levels (Fig 5C). This suggests that mRNA significant features may be standalone breast cancer predictors, while DNA methylation loci features must be considered in interaction with others to obtain highly predictive features (Table 2). This finding seems logical in the context of the regulatory functions of DNA methylation. The top n=10 of 590 significant mRNA genes from MCFS-ID ranking were verified and confirmed (based on the literature) to have a meaningful impact on cancerogenesis which implies that the choice of implemented feature selection approach is meaningful. In the METABRIC cohort [103] authors reported significant enrichment of immunologically related genes, which was also confirmed in our study, showing 79 of 590 mRNA genes being related to immunological processes. Over 75% of 590 mRNAs were down-regulated in cancer samples, which suggests that DNA hypo-methylation in cancer leads to derepression of gene expression, although such a high participation of down-expressed genes in cancer is quite surprising compared to the previously described patterns [104]. Most of the mRNAs that were up-regulated in cancer samples contributed to the cell cycle and mitosis. It fits well with the cancer cells’ specificity - increased cell division and growth. However, such a generalization may lead to erroneous conclusions. We found that a group of up-regulated genes *CCNB2*, *CCNB1*, *PLK1*, and *CDK1* was associated with the Reactome pathway “Activation Of NIMA Kinases NEK9, NEK6, NEK7” (S2 Table). Up-regulation of *PLK1* in some of the breast cancer subtypes may inhibit tumor development by interfering with cytokinesis and mitosis [105,106] as well as one of the NIMA Kinases *NEK9*, which is associated with tumor growth prevention, when upregulated [107]. Additionally, a group of genes from cluster 5, identified with the NLP clustering analysis (Table 3), showed a link with tumor suppressors, e.g. *APC* gene that, through association with other proteins, prevents the uncontrolled growth of cells [108]. Concluding, depending on the biological context, up-regulated genes may result in activation as well as repression of processes contributing to cancer development.

As mentioned before, the majority of significant mRNAs (n=590) were down-regulated in cancer cells. Pathway analysis showed that they can perturb the regulation of a wide range of Reactome metabolic pathways including ethanol oxidation [109], lipid metabolism [110,111], which are linked to breast cancer development, and also retinoid acid or neurotransmission, specifically connected with breast cancer treatment methods [112–115]. In addition to the results from the Reactome database, the NLP clustering approach unveiled a very interesting group of genes down-regulated in cancer (cluster 3, Table 3) associated with G protein-coupled receptors (GPCRs), which are cell surface receptors that detect ligands outside of the cell and initiate cellular response [116]. Moreover, in pathological states, GPCRs are over-expressed and activated in an aberrant way. This may imply certain aspects of cancer, including growth, invasion, migration, angiogenesis and metastasis [117,118]. In breast cancer multiple specific GPCRs were confirmed to participate in a plethora of autocrine and paracrine physiological effects or through activation of various ligands modulate cellular functions, which was associated with mRNA gene over-expression (revised in [119,120]). Interestingly, the genes from cluster 3 related to GPCRs were down-regulated, which is in contrast to known studies and that inconsistency should be tested *in vivo*. Another group of genes (cluster 2 Table 3) was described by keywords related to the ATP-binding cassette transporters (ABC transporters) of which some are used in ion transport that is important for muscle contraction and cardiac processes. The majority of genes in this group were down-regulated. There is evidence in the literature that ABC transporters and related to functions returned by NLP as single terms: ‘ions’, ‘transport’, ‘muscle’, ‘contraction’ and ‘cardiac’ are known to be linked with cancer biology [121]. For the ABC transporters family, both decreased and increased expression levels may be unfavorable for cancer patients, and some of the genes from this family are already molecular targets for anticancer drugs [122,123]. In this study, their levels were decreased in cancer samples, and there are reports that this may contribute to the occurrence of a more aggressive form of cancer, which was shown in *TMPRSS2-ERG-*negative prostate cancer [124]. For decreased expression of the *ABCA9* gene in epithelial ovarian cancer, a significantly shorter time to progression was observed [125]. Interestingly, in breast cancer patients the reduced *ABCA8* expression lowers 5-year patients survival rate, is present in older patients (>60 age) as well as in the three breast cancer subtypes: ER-negative, PR-negative, HER-positive [126]. Among the keywords describing the remaining clusters of genes (Table 3), both general and very specific phrases were present. It seems that NLP based clustering analysis allows for efficient linking of even small gene groups with their related processes, which we find as a big advantage. The obtained clustering results allow for a precise selection of genes having a general role from once having specific functions Further development of this NLP-based gene ontology analysis seems promising, especially as NLP is already widely and successfully used in other fields e.g. supports disease diagnosis [127].

In breast cancer samples, DNA hyper-methylation in the regulatory regions of tumor suppressor genes and hypo-methylation of oncogenes has been shown [128]. In this study MCFS-ID returned a ranking of 2006 DMSs that were significant predictors of breast cancer. They were located noticeably more often in CpG Islands (CpGI) and open seas and less often in shores than would be expected by the cytosines distribution in the entire Illumina panel. This result shows that the chance for a change in the methylation level between the tumor and the control samples depends on the genomic location. Overall, a small fraction of the returned significant DMSs was hypo-methylated in cancer samples (Fig 5C) while the majority was hyper-methylated. DMSs located in CpGIs were the most differentiated between normal and tumor samples in terms of methylation level (Fig 5B). This result corresponds very well to the pattern of generally dominant DNA hyper-methylation in breast cancer and only local disorders of DNA hypo-methylation (revised in [129]).

There were discovered putative functional associations between 59 DMS-mRNA pairs, out of which over 90% of DMS were located in distal genomic regulatory regions. DNA methylation changes of these loci may potentially affect the activity of enhancer-like regions, influencing the target-gene expression. Out of 34 genes under such enhancer-like regulatory effect, only two *KIFC1* and *HN1L* were over-expressed in cancer, both with significant negative correlation with DNA methylation. Up-regulation of *KIFC1* expression is well known for breast cancer [130] and the protein has been suggested as a chemotherapy target [131], while over-expression of *HN1L* is related to tumor invasion in breast cancer [132]. Next, based on the discovered association among DNA methylation sites within the promoter of *NKAPL* gene and TFBS of NRF1 TF shown in this study, we are confident about the presence of putative functional dependency of these molecular elements. In the proposed regulatory model, 5 hyper-methylated cytosines inhibit NRF1 binding within the *NKAPL* promoter resulting in its down-expression in tumor samples. Competition between DNA methylation and NRF1 binding to DNA sequences was confirmed [133], supporting the proposed regulatory model of *NKAPL*. Moreover, based on the results from the extensive use of MCFS-ID, it was possible to select 23 cytosines significantly associated with *NKAPL* expression (r=0.81). Among all of them only one was located within *NKAPL* promoter (cg18675097) and all the others were in the range from 1369587 to 130556755 base pairs from *NKAPL* TSS, suggesting their presence in distal regulatory regions of *NKAPL*. By using this approach we additionally selected 66 mRNA genes, whose expression was well predicted by DNA methylation and 39 of them by miRNA expression as well (see results section “Detection of miRNA and DNA methylation loci significant in the context of predicting mRNA expression levels”).

To review the significant set of n=105 miRNA returned by MCFS-ID, the analysis of associations between miRNA to mRNA was conducted resulting in the list of proteins targeted by drugs used for breast cancer treatment. For example, palbociclib (DB09073), a drug for treating metastatic breast cancer which targets proteins encoded by genes such as *CCNA2*, *CCND1*, *CDC25A*, *CDK1*, *CHEK1*, *ESR1*, *KRAS* and *PLK1* has a DScore of 0.99 while the GScore of the genes is 0.85. At the same time, other drugs with high DScore, used in breast cancer treatment, eg. ribociclib [81], abemaciclib [82], tamoxifen [134] etc. were found to be linked to miRNA reported as significant in this study (S1C Fig).

Methylated/unmethylated nucleotides within the TFBS may disturb TF binding to DNA sequence. This may change TF binding affinity, and shift the factor binding site, resulting in alternative protein complex formation, binding prevention or other TFs binding to such locus [135,136]. At the same time, it is not clear what the order of these events is, what initiates the process and what is the result [24]. In this study we identified many more TF motifs overlapping hyper-methylated DMSs than hypo-methylated, which reflects the much higher frequency of hyper-methylated cytosines among the 2006 DMSs. Moreover, almost all motifs, except EPAS1, containing hypo-methylated cytosines were identical to the TF motifs containing hyper-methylated cytosines. There are reports indicating that EPAS1 may support proliferation, migration and increase tumor cell invasiveness [137,138]. Genes encoding TFs that bind to motifs identified in sequences containing hyper-methylated cytosines (Fig 8D) belonged to, among others, cell cycle signaling pathways, transcription misregulation in cancer, the TGF-β pathway, cellular senescence, and several cancer types, including breast cancer q-value=0.0000234 (Fig 8D and S10 Table).

In the second MCFS-ID experiment, the association between DMSs and expression of genes encoding TFs, whose motifs overlapped these DMSs, was exposed. There were 7 genes encoding TFs: *MXI1*, *EPAS1*, *PLAGL1*, *E2F1*, *NR0B1*, *BHLHE41* and *ARNT2*. *PLAGL1* has been reported as a possible epigenetically regulated tumor suppressor gene [139] and *NR0B1* (also known as *DAX-1*) has been repeatedly indicated as a potential target for anti-cancer therapy in breast cancer patients [140,141]. Likewise, there is evidence that low expression of *BHLHE41*, which was also observed in the results of this study, promotes breast cancer tumor invasion [142]. The other four *EPAS1*, *MXI1*, *ARNT2* and *E2F1* are connected with the signaling pathways involved in the processes of tumor formation and development [143].

Using the epigenetic variables that functionally interact with each other, here DMSs and TFs, together with MCFS-ID and linear regression models allowed for the identification of mRNA target genes under probable epigenetic regulation (see results Fig 9A, S12 Table). Among nine target genes, four were confirmed to have linear models with high goodness of fit: *TMEM220*, *NKAPL*, *SHE*, *TGFBR3.* These genes were down-regulated in tumor samples and seem to be tumor suppressors whose activity can be regulated by DNA methylation located within TFBS of specific TFs.

Hyper-methylated DMSs may reduce these four TFs binding affinity and change gene expression. Moreover, these DMSs are located in heterochromatin regions, known to contribute to gene silencing. *NKAPL* (NKAP-like) is a cell-specific transcriptional suppressor in Notch signaling [144] and its reduced expression in cancer has been indicated by several articles, same as the relationship between DMS hyper-methylation and demonstrated change in *NKAPL* expression [93,145]. Transforming growth factor beta receptor III encoded by *TGFBR3,* binds inhibin and can mediate functional antagonism of activin signaling [146]. Decreased expression of *TGFBR3* (former *ETDL1*) causes a decreased TβRIII expression in tumor tissue resulting in tumor progression because of increased invasiveness, angiogenesis, and chance of metastasis [147,148]. Next, the SH2 (SHE) domain-containing adapter protein E possesses the Src homology 2 (SH2) domain identified in the oncoproteins Src and Fps. It functions as a regulatory module of intracellular signaling cascades by interacting with phosphotyrosine-containing target peptides [149]. The Transmembrane protein 220 (*TMEM220*) is involved in the FOXO and PI3K-Akt pathways [150] and promotes regeneration [151]. Down-expression of *TMEM220* and *SHE* genes (also in connection with hyper-methylation) has been repeatedly indicated as a significant factor important in the formation and development of cancer, but not necessarily breast cancer [62,150,152; Supplement Table S1 153]. Our results would therefore be the first to indicate the impact of these two genes in breast cancer development.

To extend the analysis of functional importance of the four genes the gene-gene interactions were used to build the network of 17 new genes connected to the initial four (Fig 9B). The functional investigation of genes with the highest number of connections (*MYC*, *NOG, TGFB1*, *VIRMA, SRC* and *AR*) in the network showed an overrepresentation of signaling pathways related to cancer processes, mainly associated with cell cycle disorders namely: proliferation, growth, differentiation, migration or apoptosis as well as patients survival. The returned KEGG pathways were consistent with the previously discussed gene biological functions. For example, the hyperactivation of the Mitogen Activated Protein Kinase (MAPK) pathway is frequently observed in many cancers, including breast cancer. It is an oncogenic pathway, and at the same time, it is crucial for the signal transduction of the ErbB protein family [154]. Some proteins from the ErbB family are oncogenes associated with proliferation and apoptosis. They are also related to cancer treatment resistance in some breast cancer subtypes [155]. At the same time, the MAPK is associated with PI3K-AKT-mTOR, i.e. the pathway which is directly related to *TGFB1* and *MYC* and indirectly with *TMEM220* and *AR*. PI3K-AKT-mTOR is associated with the processes of oncogenesis and breast cancer development, and many inhibitors of this pathway are currently in clinical trials [156,157]. Another overrepresented pathway was Hippo, which is linked with proliferation, migration and apoptosis [158] and metastasis changes [159]. The Hippo pathway is well-known for having an impact on the transforming growth factor beta (TGF-β) signaling pathway through which they may control tumor development [160,161]. Additionally, components of the TGF-β pathway play a significant role in the proliferation, cell growth and differentiation of cells, but also affect the immune system, enabling the repair or development of ongoing processes that were shown to negatively affect the patient’s condition [160,162,163]. Members of the TGF-β protein family play an essential role in apoptosis and migration, the regulation of which can have a vast impact on breast tumor development, especially at its later stage [160,163,164]. Additionally, one of the miRNA genes (hsa-miR-211) that was pointed out as significant in the presented regulatory network (Fig 9A, S5 Table), is known to participate in the TGF-β pathway [165]. The hsa-miR-211 is involved in the regulation of proliferation, migration, invasion, apoptosis and drug resistance [166] and we discovered its association with the *SHE* gene. Alterations in the expression level of hsa-miR-211 have been repeatedly reported in the context of various cancer types, but the direction of its expression level changes depending on the type of cancer. *SHE* was confirmed as an oncogene and/or tumor suppressor depending on cancer type [167]. In breast cancer, change in *SHE* expression, no matter if decreased or increased, results in metastasis and poor prognosis [166]. We are convinced that the analysis of hsa-miR-211 with the *SHE*, whose role is little known, should become the subject of detailed studies. Additionally, one of the terms related to the obtained gene-gene network was “chemical carcinogenesis”, which is connected to many environmental and chemical factors having a strong impact on the oncogenic processes including DNA methylation. Therefore, there are multiple indirect confirmations that the created network demonstrates interplay among detected epigenetic disorders, which in turn lead to subsequent changes affecting target gene expression and in turn result in disease development.

From the massive MCFS-ID computational approach, we discovered an association between cg23866403 loci and *FXYD1* and, to verify that functional putative association, we built a chromatin spatial model, using the 3D-GNOME approach. Based on the obtained results (Fig 10), we hypothesize that, in normal tissue, the lower level of cg23866403 loci methylation in the *FXYD1* gene promoter results in a shortened spatial distance to the gene enhancer and, as a result, *FXYD1* increased expression compared to cancer tissue. The significant impact of DNA methylation on chromatin structure in cancer is well studied [168], showing the appearance of changes for example due to CTCF binding disruption [169]. Moreover, the length of the DNA loops may change depending on the cohesin presence in gene expression regulatory machinery as well as the presence of CTCF in one or both anchors of a loop [170]. The relationship between distance change and gene expression change is very poorly understood and the only example we found was in *Drosophila* [171]. Based on our result, one could suggest that the change in DNA methylation affected protein binding and in consequence changed the length of the loop. Through the results presented in this study, the *NKAPL* gene was found to appear in multiple contexts making it an interesting target, especially in its transcription relation due to altered DNA methylation. Therefore, we built spatial models for it using two different experimental chromosome conformation capture protocols-ChIA-PET and PCHi-C (Fig 10). The first model allowed us to observe the change in loop length and discover that cohesin-mediated loops surrounding the *NKAPL* gene were longer in normal cell line (hTERT-HME1) than in cancerous (MCF-7), and also that loops anchored by enhancers are present only in the normal cell line. Thanks to the second model, it was noticed that in normal cell lines (MCF-10A) around the *NKAPL* gene, several times more interactions can be identified compared to the cancer cell line (MCF-7) with a simultaneous higher level of DNA methylation in cancer. Based on the spatial models obtained by both methods, we hypothesize that the reduced level of methylation in normal cells results in the formation of a much larger number of stable interactions, which translates into a more condensed chromatin region. However, this increased level of chromatin condensation in normal cell line cannot be interpreted as closed chromatin, preventing transcription and expression regulation processes. It is tighter because of the higher number of established connections. In contrast to the cancer cell line, where fewer connections result in looser chromatin structure, putatively unstable and due to that, exposed to unexpected transcriptional changes that may result in the development of potentially oncogenic changes. Chromatin organization has a significant impact on organism functioning [172], and its disruption may support pathogenic processes, e.g. through chromosomal instability which intensifies deregulation of gene expression [173]. Changes in DNA looping may cause errors in gene regulation both during cancer initiation and development [40,174]. Both the absence of such loops and the change in chromatin interaction frequency in cell lines was demonstrated to have connection with carcinogenesis [58]. Moreover, more loosely packed chromatin may reduce nuclear stiffness and at the same time increase chromatin mobility [175, 176]. This may cause chromosomal translocations and changes in transcription landscape which may contribute to oncogenic events [173,177]. This may explain why we observed more loosely packed chromatin in the cancer cell line related to the *NKAPL* gene promoter. Cohesin-mediated chromatin structures are known as regulators of Epithelial-Mesenchymal Transition (EMT) related genes, whose expression changes influence cancer progression, including breast cancer [178]. EMT may also be influenced by disturbances in the TGF-β signaling pathway, which may suggest the multilayer nature of the cohesin-mediated chromatin structures disorder [178,179]. Finally, it has to be noticed that we used TCGA tissue samples and they represent various cell heterogeneity. This may influence the study, because some of the observed values might be averaged and not confirmed as significant. At the same time, we are sure that presented here altered transcriptomic and epigenetic signals are of great value and studied further at the level of specific cell populations will bring detailed insight into gene expression regulation during breast cancer development.

## Summary

Our studies have allowed us to extract a very rich set of molecular features including mRNA expression, miRNA expression, DNA methylation allowing for a precise classification of breast cancer vs normal samples. We are certain that among them are new targets for further functional research in the context of breast cancer development and drug studies. In addition to known features validating our methodology, we have listed new candidates, and for some of them we presented possible mechanisms of their regulation. The most interesting seems to be the proposed regulatory model of *NKAPL*, *PITX1* and *TMEM220*. The computational results of 3D chromatin structure models deserve attention, which in our opinion are very interesting candidates for further studies.

## Acknowledgements

We would like to thank Małgorzata Perycz for the revision of the manuscript draft. MW, AA, KS, SG, DP research was funded by Warsaw University of Technology within the Excellence Initiative: Research University (IDUB) programme and co-supported by Polish National Science Centre (2020/37/B/NZ2/03757). Computations were performed thanks to the Laboratory of Bioinformatics and Computational Genomics, Faculty of Mathematics and Information Science, Warsaw University of Technology using Artificial Intelligence HPC platform financed by Polish Ministry of Science and Higher Education (decision no. 7054/IA/SP/2020 of 2020-08-28). The work was co-supported by National Institute of Health USA 4DNucleome grant 1U54DK107967-01 “Nucleome Positioning System for Spatiotemporal Genome Organization and Regulation”.

## Bibliography

1. Sung H, Ferlay J, Siegel RL, Laversanne M, Soerjomataram I, Jemal A, et al. Global cancer statistics 2020: GLOBOCAN estimates of incidence and mortality worldwide for 36 cancers in 185 Countries. CA: a Cancer Journal for Clinicians. 2021 Feb 4;71(3):209–49. 10.3322/caac.21660

2. Dumbrava EI, Meric-Bernstam F. Personalized cancer therapy—leveraging a knowledge base for clinical decision-making. Cold Spring Harb Mol Case Stud. 2018 Apr 1;4(2). 10.1101/mcs.a001578

3. Chopra S, Khosla M, Vidya R. Innovations and challenges in breast cancer care: a review. Medicina. 2023 May 16;59(5):957–7. 10.3390/medicina59050957

4. Bean GR, Lin CY. Breast neuroendocrine neoplasms: practical applications and continuing challenges in the era of the 5th edition of the WHO classification of breast tumours. Diagnostic Histopathology. 2021 Jan;27(4):139–47. 10.1016/j.mpdhp.2021.01.001

5. Cree IA, White VA, Indave BI, Lokuhetty D. Revising the WHO classification: female genital tract tumours. Histopathology. 2019 Dec 17;76(1):151–6. 10.1111/his.13977.

6. Blumer A, Ehrenfeucht A, Haussler D, Warmuth MK. Learnability and the Vapnik-Chervonenkis dimension. Journal of the ACM. 1989 Oct 1;36(4):929–65. 10.1145/76359.76371

7. Dramiński M, Koronacki J. rmcfs: An R Package for Monte Carlo Feature Selection and Interdependency Discovery. Journal of Statistical Software. 2018;85(12). 10.18637/jss.v085.i12

8. Dramiński M, Da̧browski MJ, Diamanti K, Koronacki J, Komorowski J, Discovering networks of interdependent features in high-dimensional problems. In Japkowicz N, Stefanowski J, editors. Big Data Analysis: New Algorithms for a New Society. Studies in Big Data, Springer, Cham; 2015. pp. 285–304.

9. Chen L, Zhou X, Zeng T, Pan X, Zhang YH, Huang T, et al. Recognizing pattern and rule of mutation signatures corresponding to cancer types. Frontiers in Cell and Developmental Biology. 2021 Aug 26;9. 10.3389/fcell.2021.712931

10. Li J, Xu Q, Wu M, Huang T, Wang Y. Pan-cancer classification based on self-normalizing neural networks and feature selection. Front Bioeng Biotechnol. 2020 Aug 4;8. 10.3389/fbioe.2020.00766

11. Li Z, Mei Z, Ding S, Chen L, Li H, Feng K, et al. Identifying methylation signatures and rules for COVID-19 with machine learning methods. Frontiers in Molecular Biosciences. 2022 May 10;9. 10.3389/fmolb.2022.908080

12. Chen L, Zhang S, Pan X, Hu X, Zhang YH, Yuan F, et al. HIV infection alters the human epigenetic landscape. Gene Therapy. 2018 Nov 15;26(1-2):29–39. 10.1038/s41434-018-0051-6

13. Li D, Lin H, Li L. Multiple feature selection strategies identified novel cardiac gene expression signature for heart failure. Frontiers in Physiology. 2020 Nov 11;11. 10.3389/fphys.2020.604241

14. Paratala BS, Dolfi SC, Khiabanian H, Rodríguez-Rodríguez L, Ganesan S, Hirshfield KM. Emerging role of genomic rearrangements in breast cancer: applying knowledge from other cancers. Biomarkers in cancer. 2016 Jan 1;8:1–14. 10.4137/bic.s34417

15. Banerji S, Cibulskis K, Rangel-Escareno C, Brown KK, Carter SL, Frederick AM, et al. Sequence analysis of mutations and translocations across breast cancer subtypes. Nature. 2012 Jun 1;486(7403):405–9. 10.1038/nature11154

16. Kagohara LT, Stein-O’Brien GL, Kelley D, Flam E, Wick HC, Danilova LV, et al. Epigenetic regulation of gene expression in cancer: techniques, resources and analysis. Briefings in Functional Genomics. 2017 Aug 11;17(1):49–63. 10.1093/bfgp/elx018

17. Dabrowski MJ, Draminski M, Diamanti K, Stepniak K, Mozolewska MA, Teisseyre P, et al. Unveiling new interdependencies between significant DNA methylation sites, gene expression profiles and glioma patients survival. Scientific Reports. 2018 Mar 13;8(1). 10.1038/s41598-018-22829-1

18. Cao J, Yan Q. Cancer epigenetics, tumor immunity, and immunotherapy. Trends in Cancer. 2020 Mar;6(7):580–92. 10.1016/j.trecan.2020.02.003

19. Zhong Q, Fan J, Chu H, Pang M, Li J, Fan Y, et al. Integrative analysis of genomic and epigenetic regulation of endometrial cancer. Aging. 2020 May 15;12(10):9260–74. 10.18632/aging.103202

20. Vezzani B, Carinci M, Previati M, Giacovazzi S, Della Sala M, Gafà R, et al. Epigenetic regulation: a link between inflammation and carcinogenesis. Cancers. 2022 Jan 1;14(5):1221. 10.3390/cancers14051221

21. Portela A, Esteller M. Epigenetic modifications and human disease. Nature Biotechnology. 2010 Oct;28(10):1057–68. 10.1038/nbt.1685

22. Skvortsova K, Stirzaker C, Taberlay P. The DNA methylation landscape in cancer. Essays in Biochemistry. 2019 Dec 1;63(6):797–811. 10.1042/EBC20190037

23. Yan W, Herman JG, Guo M. Epigenome-based personalized medicine in human cancer. Epigenomics. 2016 Jan;8(1):119–33. 10.2217/epi.15.84

24. Héberlé É, Bardet A. Sensitivity of transcription factors to DNA methylation. Essays in Biochemistry. 2019 Nov 22;63(6):727–41. 10.1042/ebc20190033

25. Blattler A, Farnham PJ. Cross-talk between site-specific transcription factors and DNA methylation states. J Biol Chem. 2013 Nov;288(48):34287–94. 10.1074/jbc.r113.512517

26. Costa FF, Paixão VA, Cavalher FP, Ribeiro KB, Cunha IW, Rinck Jr JA, et al. SATR-1 hypomethylation is a common and early event in breast cancer. Cancer genetics and cytogenetics. 2006 Mar 1;165(2):135–43. 10.1016/j.cancergencyto.2005.07.023

27. Yi J, Gao R, Chen Y, Yang Z, Han P, Zhang H, et al. Overexpression of NSUN2 by DNA hypomethylation is associated with metastatic progression in human breast cancer. Oncotarget. 2016 Nov 30;8(13):20751–65. 10.18632/oncotarget.10612

28. Choi JY, James SR, Link PA, McCann SE, Hong CC, Davis W, et al. Association between global DNA hypomethylation in leukocytes and risk of breast cancer. Carcinogenesis. 2009 Jul 7;30(11):1889–97. 10.1093/carcin/bgp143

29. Martin TA, Goyal A, Watkins G, Jiang WG. Expression of the transcription factors Snail, Slug, and Twist and their clinical significance in human breast cancer. Annals of Surgical Oncology. 2005 Apr 19;12(6):488–96. 10.1245/aso.2005.04.010

30. Dulaimi E, Hillinck J, Ibanez de Caceres I, Al-Saleem T, Cairns P. Tumor suppressor gene promoter hypermethylation in serum of breast cancer patients. Clinical Cancer Research. 2004 Sep 15;10(18):6189–93. 10.1158/1078-0432.ccr-04-0597

31. Alvarez C, Tapia T, Cornejo V, Fernandez W, Muñoz A, Camus M, et al. Silencing of tumor suppressor genes RASSF1A, SLIT2, and WIF1 by promoter hypermethylation in hereditary breast cancer. Molecular Carcinogenesis. 2012 Feb 7;52(6):475–87. 10.1002/mc.21881

32. Su J, Huang YH, Cui X, Wang X, Zhang X, Lei Y, et al. Homeobox oncogene activation by pan-cancer DNA hypermethylation. Genome Biology. 2018 Aug 10;19(108). 10.1186/s13059-018-1492-3

33. Spainhour JC, Lim HS, Yi SV, Qiu P. Correlation patterns between DNA methylation and gene expression in the cancer genome atlas. Cancer Informatics. 2019 Jan;18:1–11. 10.1177/1176935119828776

34. Yao Q, Chen Y, Zhou X. The roles of microRNAs in epigenetic regulation. Current Opinion in Chemical Biology. 2019 Aug;51:11–7. 10.1016/j.cbpa.2019.01.024

35. Ying SY, Chang DC, Lin SL. The microRNA (miRNA): Overview of the RNA genes that modulate gene function. Molecular Biotechnology. 2008;38(3):257–68. 10.1007/s12033-007-9013-8

36. Pavlíková L, Šereš M, Breier A, Sulová Z. The roles of microRNAs in cancer multidrug resistance. Cancers. 2022 Feb 21;14(4):1090–0. 10.3390/cancers14041090

37. Muñoz JP, Pérez-Moreno P, Pérez Y, Calaf GM. The role of microRNAs in breast cancer and the challenges of their clinical application. Diagnostics. 2023 Jan 1;13(19):3072. 10.3390/diagnostics13193072

38. Umer HM, Cavalli M, Dabrowski M, Diamanti K, Kruczyk M, Pan G, et al. A significant regulatory mutation burden at a high-affinity position of the CTCF motif in gastrointestinal cancers. Human Mutation. 2016 Jun 2;37(9):904–13. 10.1002/humu.23014

39. Monteagudo-Sánchez A, Noordermeer D, Greenberg MVC. The impact of DNA methylation on CTCF-mediated 3D genome organization. Nature Structural & Molecular Biology. 2024 Mar;31(3):404–12. 10.1038/s41594-024-01241-6

40. Grabowicz IE, Wilczyński B, Kamińska B, Roura AJ, Wojtaś B, Dąbrowski MJ. The role of epigenetic modifications, long-range contacts, enhancers and topologically associating domains in the regulation of glioma grade-specific genes. Scientific Reports. 2021 Aug 2;11(1). 10.1038/s41598-021-95009-3

41. Poulos RC, Sloane MA, Hesson LB, Wong JWH. The search for cis-regulatory driver mutations in cancer genomes. Oncotarget. 2015 Aug 20;6(32):32509–25. 10.18632/oncotarget.5085

42. Draminski M, Rada-Iglesias A, Enroth S, Wadelius C, Koronacki J, Komorowski J. Monte Carlo feature selection for supervised classification. Bioinformatics. 2007 Nov 28;24(1):110–7. 10.1093/bioinformatics/btm486

43. Kuleshov MV, Jones MR, Rouillard AD, Fernandez NF, Duan Q, Wang Z, et al. Enrichr: a comprehensive gene set enrichment analysis web server 2016 update. Nucleic Acids Research. 2016 May 3;44(W1):W90–7. 10.1093/nar/gkw377

44. Hinrichs AS, Karolchik D, Baertsch R, Barber GP, Bejerano G, Clawson H, et al. The UCSC Genome Browser Database: update 2006. Nucleic Acids Research. 2006 Jan 1;34:D590–8. 10.1093/nar/gkj144

45. Quinlan AR, Hall IM. BEDTools: a flexible suite of utilities for comparing genomic features. Bioinformatics. 2010 Jan 28;26(6):841–2. 10.1093/bioinformatics/btq033

46. Weisenberger DJ, Van Den Berg D, Pan F, Berman BP, Laird PW. Comprehensive DNA methylation analysis on the Illumina ® Infinium ® Assay Platform [Internet]. www.illumina.com. 2008. Available from: https://www.illumina.com/content/dam/illumina-marketing/documents/products/appnotes/appnote_dna_methylation_analysis_infinium.pdf

47. Chen Y, Wang X. miRDB: an online database for prediction of functional microRNA targets. Nucleic Acids Research. 2020 Jan 8;48(D1):D127–31. 10.1093/nar/gkz757

48. Szklarczyk D, Kirsch R, Koutrouli M, Nastou K, Mehryary F, Hachilif R, et al. The STRING database in 2023: protein–protein association networks and functional enrichment analyses for any sequenced genome of interest. Nucleic Acids Research. 2022 Nov 12;51(D1):D638–46. 10.1093/nar/gkac1000

49. Pian C, Zhang G, Gao L, Fan X, Li F. miR+Pathway: the integration and visualization of miRNA and KEGG pathways. Briefings in Bioinformatics. 2019 Jan 16;21(2):699–708. 10.1093/bib/bby128

50. Pongubala JMR, Murre C. Spatial organization of chromatin: transcriptional control of adaptive immune cell development. Frontiers in Immunology. 2021 Mar 29;12. 10.3389/fimmu.2021.633825

51. Kulakovskiy IV, Vorontsov IE, Yevshin IS, Sharipov RN, Fedorova AD, Rumynskiy EI, et al. HOCOMOCO: towards a complete collection of transcription factor binding models for human and mouse via large-scale ChIP-Seq analysis. Nucleic Acids Research. 2017 Nov 11;46(D1):D252–9. 10.1093/nar/gkx1106

52. Stojnic R, Diez D. PWMEnrich: PWM enrichment analysis. Bioconductor. 2024. Available from: https://bioconductor.org/packages/release/bioc/html/PWMEnrich.html

53. Grant CE, Bailey TL, Noble WS. FIMO: scanning for occurrences of a given motif. Bioinformatics. 2011 Feb 16;27(7):1017–8. 10.1093/bioinformatics/btr064

54. Mahony S, Benos PV. STAMP: a web tool for exploring DNA-binding motif similarities. Nucleic Acids Research. 2007 May 8;35:W253–8. 10.1093/nar/gkm272

55. Rodchenkov I, Babur O, Luna A, Aksoy BA, Wong JV, Fong D, et al. Pathway Commons 2019 Update: integration, analysis and exploration of pathway data. Nucleic Acids Research. 2019 Oct 24;48(D1):D489–97. 10.1093/nar/gkz946

56. Szałaj P, Tang Z, Michalski P, Pietal MJ, Luo OJ, Sadowski M, et al. An integrated 3-Dimensional Genome Modeling Engine for data-driven simulation of spatial genome organization. Genome Research. 2016 Oct 27;26(12):1697–709. 10.1101/gr.205062.116

57. Kadlof M, Rozycka J, Dariusz Plewczynski. Spring Model – chromatin modeling tool based on OpenMM. Methods. 2019 Nov 30;181-182:62–9. 10.1016/j.ymeth.2019.11.014

58. Grubert F, Srivas R, Spacek D, Kasowski M, Ruiz-Velasco M, Sinnott-Armstrong N, et al. Landscape of cohesin-mediated chromatin loops in the human genome. Nature. 2020 Jul 29;583(7818):737–43. 10.1038/s41586-020-2151-x

59. Beesley J, Sivakumaran H, Moradi Marjaneh M, Lima LG, Hillman KM, Kaufmann S, et al. Chromatin interactome mapping at 139 independent breast cancer risk signals. Genome Biology. 2020 Jan 7;21(1). 10.1186/s13059-019-1877-y

60. Cestarelli V, Fiscon G, Felici G, Bertolazzi P, Weitschek E. CAMUR: Knowledge extraction from RNA-seq cancer data through equivalent classification rules. Bioinformatics. 2015 Oct 30;32(5):697–704. 10.1093/bioinformatics/btv635

61. Huang J, Sun Y, Chen H, Liao Y, Li S, Chen C, et al. ADAMTS5 acts as a tumor suppressor by inhibiting migration, invasion and angiogenesis in human gastric cancer. Gastric Cancer. 2018 Aug 13;22(2):287–301. 10.1007/s10120-018-0866-2

62. Choi B, Han TS, Min J, Hur K, Lee SM, Lee HJ, et al. MAL and TMEM220 are novel DNA methylation markers in human gastric cancer. Biomarkers. 2016 Sep 22;22(1):35–44. 10.1080/1354750x.2016.1201542

63. Liu G, Li J, Zhang CY, Huang DY, Xu JW. ARHGAP20 Expression inhibited HCC progression by regulating the PI3K-AKT signaling pathway. Journal of Hepatocellular Carcinoma. 2021 Apr;8:271–84. 10.2147/jhc.s298554

64. Bai Y, Wei C, Zhong Y, Zhang Y, Long J, Huang S, et al. Development and validation of a prognostic nomogram for gastric cancer based on DNA methylation-driven differentially expressed genes. International Journal of Biological Sciences. 2020;16(7):1153–65. 10.7150/ijbs.41587

65. Giussani M, Landoni E, Merlino G, Turdo F, Veneroni S, Paolini B, et al. Extracellular matrix proteins as diagnostic markers of breast carcinoma. Journal of Cellular Physiology. 2018 Mar 9;233(8):6280–90. 10.1002/jcp.26513

66. Zhuang Y, Li X, Zhan P, Pi G, Wen G. MMP11 promotes the proliferation and progression of breast cancer through stabilizing Smad2 protein. Oncology Reports. 2021 Feb 3;45(4). 10.3892/or.2021.7967

67. Ozturk S, Papageorgis P, Wong CK, Lambert AW, Abdolmaleky HM, Thiagalingam A, et al. SDPR functions as a metastasis suppressor in breast cancer by promoting apoptosis. Proceedings of the National Academy of Sciences. 2016 Jan 6;113(3):638–43. 10.1073/pnas.1514663113

68. Nema R, Kumar A. Sphingosine-1-phosphate catabolizing enzymes predict better prognosis in triple-negative breast cancer patients and correlates with tumor-infiltrating immune cells. Frontiers in Molecular Biosciences. 2021 Jun 21;8. 10.3389/fmolb.2021.697922

69. Zhang W, Wang H, Qi Y, Li S, Geng C. Epigenetic study of early breast cancer (EBC) based on DNA methylation and gene integration analysis. Scientific Reports. 2022 Feb 7;12(1). 10.1038/s41598-022-05486-3

70. Bao Y, Wang L, Shi L, Yun F, Liu X, Chen Y, et al. Transcriptome profiling revealed multiple genes and ECM-receptor interaction pathways that may be associated with breast cancer. Cellular & Molecular Biology Letters. 2019 Jun 6;24(1). 10.1186/s11658-019-0162-0

71. Gillespie M, Jassal B, Stephan R, Milacic M, Rothfels K, Senff-Ribeiro A, et al. The reactome pathway knowledgebase 2022. Nucleic Acids Research. 2021 Nov 12;50(D1):D687–92. 10.1093/nar/gkab1028

72. Liu C, Pan H, Torng P, Fan M, Mao T. SRPX and HMCN1 regulate cancer-associated fibroblasts to promote the invasiveness of ovarian carcinoma. Oncology Reports. 2019 Oct 17;42:2706–15. 10.3892/or.2019.7379

73. Hilwi M, Shulman K, Naroditsky I, Feld S, Gross-Cohen M, Boyango I, et al. Nuclear localization of heparanase 2 (Hpa2) attenuates breast carcinoma growth and metastasis. Cell death and disease. 2024 Mar 22;15(3). 10.1038/s41419-024-06596-8

74. Kodigepalli KM, Bowers K, Sharp A, Nanjundan M. Roles and regulation of phospholipid scramblases. FEBS Letters. 2014 Dec 3;589(1):3–14. 10.1016/j.febslet.2014.11.036

75. Wan H, Gao W, Zhang W, Tao Z, Lu X, Chen F, et al. Network-based inference of master regulators in epithelial membrane protein 2-treated human RPE cells. BMC genomic data. 2022 Jul;23(1). 10.1186/s12863-022-01047-9

76. Wang P, Wu Y, Li Y, Zheng J, Tang J. A novel RING finger E3 ligase RNF186 regulate ER stress-mediated apoptosis through interaction with BNip1. Cellular Signalling. 2013 Nov;25(11):2320–33. 10.1016/j.cellsig.2013.07.016

77. Hu Y, Jiang N, Wang X, Wu X, Bo J, Chen Y, et al. Systematic pan-cancer analysis of RNF186 with potential implications in progression and prognosis in human cancer. Life sciences. 2024 Jan;338. 10.1016/j.lfs.2023.122389

78. Anstee JE, Feehan KT, Opzoomer JW, Dean I, Muller HP, Bahri M, et al. LYVE-1+ macrophages form a collaborative CCR5-dependent perivascular niche that influences chemotherapy responses in murine breast cancer. Developmental Cell. 2023 Jul 5;58(17). 10.1016/j.devcel.2023.06.006

79. Gad AA, Balenga N. The Emerging Role of Adhesion GPCRs in Cancer. ACS Pharmacology & Translational Science. 2020 Jan 13;3(1):29–42. 10.1021/acsptsci.9b00093

80. Ng PKS, Lau CPY, Lam EKY, Li SSK, Lui VWY, Yeo W, et al. Hypermethylation of NF-κB-Activating Protein-Like (NKAPL) promoter in hepatocellular carcinoma suppresses Its expression and predicts a poor prognosis. Digestive Diseases and Sciences. 2018 Jan 20;63(3):676–86. 10.1007/s10620-018-4929-3

81. Hortobagyi GN, Stemmer SM, Burris HA, Yap YS, Sonke GS, Hart L, et al. Overall survival with ribociclib plus letrozole in advanced breast cancer. New England Journal of Medicine. 2022 Mar 10;386(10):942–50. 10.1056/nejmoa2114663

82. Tolaney SM, Beeram M, Beck JT, Conlin A, Dees EC, Puhalla SL, et al. Abemaciclib in combination with endocrine therapy for patients with hormone receptor-positive, HER2-negative metastatic breast cancer: a phase 1b study. Frontiers in Oncology. 2022 Feb 10;11. 10.3389/fonc.2021.810023

83. Kanehisa M, Furumichi M, Sato Y, Ishiguro-Watanabe M, Tanabe M. KEGG: integrating viruses and cellular organisms. Nucleic Acids Research. 2020 Oct 30;49(D1):D545–51. 10.1093/nar/gkaa970

84. Colak S, ten Dijke P. Targeting TGF-β Signaling in Cancer. Trends in Cancer. 2017 Jan;3(1):56–71. 10.1016/j.trecan.2016.11.008

85. Michaelis M, Doerr HW, Cinatl J. The story of human cytomegalovirus and cancer: increasing evidence and open questions. Neoplasia (New York, NY). 2009 Jan 1;11(1):1–9. 10.1593/neo.81178

86. Zhao J, Xu Y. PITX1 plays essential functions in cancer. Frontiers in Oncology. 2023 Sep 29;13. 10.3389/fonc.2023.1253238

87. Vidula N, Yau C, Wolf D, Rugo HS. Androgen receptor gene expression in primary breast cancer. npj Breast Cancer. 2019 Dec;5(1). 10.1038/s41523-019-0142-6

88. Fallah Y, Brundage J, Allegakoen P, Shajahan-Haq AN. MYC-driven pathways in breast cancer subtypes. Biomolecules. 2017 Jul 11;7(3):53. 10.3390/biom7030053

89. Pietri E, Conteduca V, Andreis D, Massa I, Melegari E, Sarti S, et al. Androgen receptor signaling pathways as a target for breast cancer treatment. Endocrine-Related Cancer. 2016 Oct;23(10):R485–98. 10.1530/ERC-16-0190

90. Hardy KM, Booth BW, Hendrix MJC, Salomon DS, Strizzi L. ErbB/EGF signaling and EMT in mammary development and breast cancer. Journal of Mammary Gland Biology and Neoplasia. 2010 Jun 1;15(2):191–9. 10.1007/s10911-010-9172-2

91. Thanasupawat T, Glogowska A, Nivedita-Krishnan S, Wilson B, Klonisch T, Hombach-Klonisch S. Emerging roles for the relaxin/RXFP1 system in cancer therapy. Molecular and Cellular Endocrinology. 2019 May 1;487:85–93. 10.1016/j.mce.2019.02.001

92. Yang S, Chen K, Cao K, Xu S, Ma C, Cai Y, et al. miR-182-5p Inhibits NKAPL expression and Promotes the Proliferation of Osteosarcoma. Biotechnology and Bioprocess Engineering. 2021 Oct 1;26(5):758–66. 10.1007/s12257-021-0019-z

93. Zhang X, Kang X, Jin L, Bai J, Zhang H, Liu W, et al. ABCC9, NKAPL, and TMEM132C are potential diagnostic and prognostic markers in triple-negative breast cancer. Cell Biology International. 2020 Jul 1;44(10):2002–10. 10.1002/cbin.11406

94. Silva R, Glennon K, Metoudi M, Moran B, Salta S, Slattery K, et al. Unveiling the epigenomic mechanisms of acquired platinum-resistance in high-grade serous ovarian cancer. International journal of cancer. 2023 Mar 20;153(1):120–32. 10.1002/ijc.34496

95. Morales-Martínez M, Vega MI. Role of MicroRNA-7 (MiR-7) in Cancer Physiopathology. International Journal of Molecular Sciences. 2022 Jan 1;23(16):9091. 10.3390/ijms23169091

96. Schneider E, Pliushch G, El Hajj N, Galetzka D, Puhl A, Schorsch M, et al. Spatial, temporal and interindividual epigenetic variation of functionally important DNA methylation patterns. Nucleic Acids Research. 2010 Mar 1;38(12):3880–90. 10.1093/nar/gkq126

97. Cedar H. DNA methylation and gene activity. Cell. 1988 Apr;53(1):3–4. 10.1016/0092-8674(88)90479-5

98. Szalaj P, Michalski PJ, Wróblewski P, Tang Z, Kadlof M, Mazzocco G, et al. 3D-GNOME: an integrated web service for structural modeling of the 3D genome. Nucleic Acids Research. 2016 May 16;44(W1):W288–93. 10.1093/nar/gkw437

99. Wlasnowolski M, Sadowski M, Czarnota T, Jodkowska K, Szalaj P, Tang Z, et al. 3D-GNOME 2.0: a three-dimensional genome modeling engine for predicting structural variation-driven alterations of chromatin spatial structure in the human genome. Nucleic Acids Research. 2020 May 22;48(W1):W170–6. 10.1093/nar/gkaa388

100. Pettersen EF, Goddard TD, Huang CC, Couch GS, Greenblatt DM, Meng EC, et al. UCSF Chimera--a visualization system for exploratory research and analysis. Journal of Computational Chemistry. 2004;25(13):1605–12. 10.1002/jcc.20084

101. Javierre BM, Burren OS, Wilder SP, Kreuzhuber R, Hill SM, Sewitz S, et al. Lineage-Specific Genome Architecture Links Enhancers and Non-coding Disease Variants to Target Gene Promoters. Cell. 2016 Nov;167(5):1369–1384.e19. 10.1016/j.cell.2016.09.037

102. Achinger-Kawecka J, Stirzaker C, Portman N, Campbell E, Chia KM, Du Q, et al. The potential of epigenetic therapy to target the 3D epigenome in endocrine-resistant breast cancer. Nature Structural & Molecular Biology. 2024 Jan 5;31(3):498–512. 10.1038/s41594-023-01181-7

103. Mei J, Zhao J, Fu Y. Molecular classification of breast cancer using the mRNA expression profiles of immune-related genes. Scientific Reports. 2020 Mar 16;10(1). 10.1038/s41598-020-61710-y

104. Deng JL, Xu Y, Wang G. Identification of potential crucial genes and key pathways in breast cancer using bioinformatic analysis. Frontiers in Genetics. 2019 Aug 2;10. 10.3389/fgene.2019.00695

105. de Cárcer G, Venkateswaran SV, Salgueiro L, El Bakkali A, Somogyi K, Rowald K, et al. Plk1 overexpression induces chromosomal instability and suppresses tumor development. Nature Communications. 2018 Aug 1;9(1):3012. 10.1038/s41467-018-05429-5

106. Lashen A, Toss MS, Wootton LL, Green A, Mongan NP, Madhusudan S, et al. Characteristics and prognostic significance of polo-like kinase-1 (PLK1) expression in breast cancer. Histopathology. 2023 May 24;83(3):414–25. 10.1111/his.14960

107. Xu Z, Shen W, Pan A, Sun F, Zhang J, Gao P, et al. Decreased Nek9 expression correlates with aggressive behaviour and predicts unfavourable prognosis in breast cancer. Pathology. 2020 Apr;52(3):329–35. 10.1016/j.pathol.2019.11.008

108. Lesko AC, Goss KH, Yang FF, Schwertner A, Hulur I, Onel K, et al. The APC tumor suppressor is required for epithelial cell polarization and three-dimensional morphogenesis. Biochimica et Biophysica Acta (BBA) - Molecular Cell Research. 2015 Mar;1853(3):711–23. 10.1016/j.bbamcr.2014.12.036

109. Seitz HK, Stickel F. Acetaldehyde as an underestimated risk factor for cancer development: role of genetics in ethanol metabolism. Genes & Nutrition. 2010 Jun 1;5(2):121–8. 10.1007/s12263-009-0154-1

110. Fu Y, Zou T, Shen X, Nelson PJ, Li J, Wu C, et al. Lipid metabolism in cancer progression and therapeutic strategies. MedComm. 2020 Dec 24;2(1):27–59. 10.1002/mco2.27

111. Long J, Zhang CJ, Zhu N, Du K, Yin YF, Tan X, et al. Lipid metabolism and carcinogenesis, cancer development. American journal of cancer research. 2018;8(5):778–91.

112. Chen MC, Hsu SL, Lin H, Yang TY. Retinoic acid and cancer treatment. BioMedicine. 2014 Nov 28;4(22). 10.7603/s40681-014-0022-1

113. Jan N, Sofi S, Qayoom H, Haq BU, Shabir AM, Mir MA. Targeting breast cancer stem cells through retinoids: a new hope for treatment. Critical Reviews in Oncology/Hematology. 2023 Dec 1;192:104156. 10.1016/j.critrevonc.2023.104156

114. Costantini L, Molinari R, Farinon B, Merendino N. Retinoic acids in the treatment of most lethal solid cancers. Journal of Clinical Medicine. 2020 Jan 28;9(2):360. 10.3390/jcm9020360

115. Li RQ, Zhao XH, Zhu Q, Liu T, Hondermarck H, Thorne RF, et al. Exploring neurotransmitters and their receptors for breast cancer prevention and treatment. Theranostics. 2023;13(3):1109–29. 10.7150/thno.81403

116. Rosenbaum DM, Rasmussen SGF, Kobilka BK. The structure and function of G-protein-coupled receptors. Nature. 2009 May;459(7245):356–63. 10.1038/nature08144

117. Dorsam RT, Gutkind JS. G-protein-coupled receptors and cancer. Nature Reviews Cancer. 2007 Feb;7(2):79–94. 10.1038/nrc2069

118. De Francesco E, Sotgia F, Clarke R, Lisanti M, Maggiolini M. G Protein-Coupled receptors at the crossroad between physiologic and pathologic angiogenesis: old paradigms and emerging concepts. International Journal of Molecular Sciences. 2017 Dec 14;18(12):2713. 10.3390/ijms18122713

119. Singh A, Nunes JJ, Ateeq B. Role and therapeutic potential of G-protein coupled receptors in breast cancer progression and metastases. European Journal of Pharmacology. 2015 Sep 1;763:178–83. 10.1016/j.ejphar.2015.05.011

120. Lappano R, Jacquot Y, Maggiolini M. GPCR modulation in breast cancer. International Journal of Molecular Sciences. 2018 Dec 2;19(12):3840. 10.3390/ijms19123840

121. He J, Fortunati E, Liu DX, Yan L. Pleiotropic roles of ABC transporters in breast cancer. International Journal of Molecular Sciences. 2021 Mar 21;22(6):3199–9. 10.3390/ijms22063199

122. Chen Y, Gera L, Zhang S, Li X, Yang Y, Mamouni K, et al. Small molecule BKM1972 inhibits human prostate cancer growth and overcomes docetaxel resistance in intraosseous models. Cancer Letters. 2019 Apr 1;446:62–72. 10.1016/j.canlet.2019.01.010

123. Muriithi W, Wanjiku Macharia L, Pilotto Heming C, Lima Echevarria J, Nyachieo A, Niemeyer Filho P, et al. ABC transporters and the hallmarks of cancer: roles in cancer aggressiveness beyond multidrug resistance. Cancer Biology and Medicine. 2020;17(2):253–69. 10.20892/j.issn.2095-3941.2019.0284

124. Demidenko R, Razanauskas D, Daniunaite K, Lazutka JR, Jankevicius F, Jarmalaite S. Frequent down-regulation of ABC transporter genes in prostate cancer. BMC Cancer. 2015 Oct 12;15(1). 10.1186/s12885-015-1689-8

125. Elsnerova K, Mohelnikova-Duchonova B, Cerovska E, Ehrlichova M, Gut I, Rob L, et al. Gene expression of membrane transporters: importance for prognosis and progression of ovarian carcinoma. Oncology Reports. 2016 Jan 28;35(4):2159–70. 10.3892/or.2016.4599

126. Lv C, Yang H, Yu J, Dai X. ABCA8 inhibits breast cancer cell proliferation by regulating the AMP activated protein kinase/mammalian target of rapamycin signaling pathway. Environmental Toxicology. 2022 Feb 22;37(6):1423–31. 10.1002/tox.23495

127. Agius R, Parviz M, Niemann CU. Artificial intelligence models in chronic lymphocytic leukemia – recommendations toward state-of-the-art. Leukemia & lymphoma. 2021 Oct 6;63(2):265–78. 10.1080/10428194.2021.1973672

128. Wilting RH, Dannenberg JH. Epigenetic mechanisms in tumorigenesis, tumor cell heterogeneity and drug resistance. Drug Resistance Updates. 2012 Feb 1;15(1-2):21–38. 10.1016/j.drup.2012.01.008

129. Ma L, Li C, Yin H, Huang J, Yu S, Zhao J, et al. The mechanism of DNA methylation and miRNA in breast cancer. International Journal of Molecular Sciences. 2023 Jan 1;24(11):9360. 10.3390/ijms24119360

130. Li Y, Lu W, Chen D, Boohaker RJ, Zhai L, Padmalayam I, et al. KIFC1 is a novel potential therapeutic target for breast cancer. Cancer Biology & Therapy. 2015 Jul 15;16(9):1316–22. 10.1080/15384047.2015.1070980

131. Xiao YX, Yang WX. KIFC1: a promising chemotherapy target for cancer treatment? Oncotarget. 2016 Apr 19;7(30):48656–70. 10.18632/oncotarget.8799

132. Jiao D, Zhang J, Chen P, Guo X, Qiao J, Zhu J, et al. HN1L promotes migration and invasion of breast cancer by up-regulating the expression of HMGB1. Journal of Cellular and Molecular Medicine. 2020 Nov 16;25(1):397–410. 10.1111/jcmm.16090

133. Domcke S, Bardet AF, Adrian Ginno P, Hartl D, Burger L, Schübeler D. Competition between DNA methylation and transcription factors determines binding of NRF1. Nature. 2015 Dec;528(7583):575–9. 10.1038/nature16462

134. Osborne CK. Tamoxifen in the treatment of breast cancer. Wood AJJ, editor. New England Journal of Medicine. 1998 Nov 26;339(22):1609–18. 10.1056/nejm199811263392207

135. Hu S, Wan J, Su Y, Song Q, Zeng Y, Nguyen HN, et al. DNA methylation presents distinct binding sites for human transcription factors. eLife. 2013 Sep 3;2. 10.7554/eLife.00726

136. Zhu H, Wang G, Qian J. Transcription factors as readers and effectors of DNA methylation. Nature Reviews Genetics. 2016 Aug 1;17(9):551–65. 10.1038/nrg.2016.83

137. Zhao J, Bai Z, Feng F, Song E, Du F, Zhao J, et al. Cross-talk between EPAS-1/HIF-2α and PXR signaling pathway regulates multi-drug resistance of stomach cancer cell. The International Journal of Biochemistry & Cell Biology. 2016 Jan 16;72:73–88. 10.1016/j.biocel.2016.01.006

138. Lu X, Zhang W, Zhang J, Ren D, Zhao P, Ying Y. EPAS1, a hypoxia- and ferroptosis-related gene, promotes malignant behaviour of cervical cancer by ceRNA and super-enhancer. Journal of Cellular and Molecular Medicine. 2024 May 1;28(9). 10.1111/jcmm.18361

139. Abdollahi A, Pisarcik DA, Roberts DD, Weinstein JK, Cairns P, Hamilton TA. LOT1 (PLAGL1/ZAC1), the candidate tumor suppressor gene at chromosome 6q24–25, is epigenetically regulated in cancer. Journal of Biological Chemistry. 2003 Feb 21;278(8):6041–9. 10.1074/jbc.m210361200

140. Lanzino M, Maris P, Sirianni R, Barone I, Casaburi I, Chimento A, et al. DAX-1, as an androgen-target gene, inhibits aromatase expression: a novel mechanism blocking estrogen-dependent breast cancer cell proliferation. Cell Death & Disease. 2013 Jul;4(7):e724. 10.1038/cddis.2013.235

141. Chae BJ, Lee A, Bae JS, Song BJ, Jung SS. Expression of nuclear receptor DAX-1 and androgen receptor in human breast cancer. Journal of Surgical Oncology. 2011 Jan 15;103(8):768–72. 10.1002/jso.21861

142. Zhang D, Zheng Q, Wang C, Zhao N, Liu Y, Wang E. BHLHE41 suppresses MCF-7 cell invasion via MAPK/JNK pathway. Journal of Cellular and Molecular Medicine. 2020 Feb 19;24(7):4001–10. 10.1111/jcmm.15033

143. Keith B, Johnson RS, Simon MC. HIF1α and HIF2α: sibling rivalry in hypoxic tumor growth and progression. Nature reviews Cancer. 2011 Dec 15;12(1):9–22. 10.1038/nrc3183

144. Okuda H, Kiuchi H, Takao T, Miyagawa Y, Tsujimura A, Nonomura N, et al. A novel transcriptional factor Nkapl is a germ cell-specific suppressor of notch signaling and is indispensable for spermatogenesis. PLoS ONE. 2015 Apr 14;10(4):e0124293–3. 10.1371/journal.pone.0124293

145. Li M, Sun Q, Wang X. Transcriptional landscape of human cancers. Oncotarget. 2017 Mar 2;8(21):34534–51. 10.18632/oncotarget.15837

146. Lewis KA, Gray PC, Blount AL, MacConell LA, Wiater E, Bilezikjian LM, et al. Betaglycan binds inhibin and can mediate functional antagonism of activin signalling. Nature. 2000 Mar 1;404(6776):411–4. 10.1038/35006129

147. Bao S, He G. Identification of key genes and key pathways in breast cancer based on machine learning. Medical Science Monitor. 2022 Apr 25;28. 10.12659/msm.935515

148. Dong M, How T, Kirkbride KC, Gordon KJ, Lee JT, Hempel N, et al. The type III TGF-β receptor suppresses breast cancer progression. Journal of Clinical Investigation. 2007 Jan 2;117(1):206–17. 10.1172/jci29293

149. Mayer BJ, Baltimore D. Signalling through SH2 and SH3 domains. Trends in Cell Biology. 1993 Jan;3(1):8–13. 10.1016/0962-8924(93)90194-6

150. Li T, Guan L, Tang G, He B, Huang L, Wang J, et al. Downregulation of TMEM220 promotes tumor progression in Hepatocellular Carcinoma. Cancer Gene Therapy. 2021 Jul 28;29(6):835–44. 10.1038/s41417-021-00370-0

151. Jung SY, Kim DY, Yune TY, Shin DH, Baek SB, Kim CJ. Treadmill exercise reduces spinal cord injury-induced apoptosis by activating the PI3K/Akt pathway in rats. Experimental and Therapeutic Medicine. 2013 Dec 17;7(3):587–93. 10.3892/etm.2013.1451

152. Su PH, Hsu YC, Huang R, Weng YC, Wang HC, Chen Y, et al. Methylomics of nitroxidative stress on precancerous cells reveals DNA methylation alteration at the transition from in situ to invasive cervical cancer. Oncotarget. 2017 Jun 6;8(39):65281–91. 10.18632/oncotarget.18370

153. Zhang X, Zhang H, Fan C, Hildesjö C, Shen B, Sun XF. Loss of CHGA protein as a potential biomarker for colon cancer diagnosis: a study on biomarker discovery by machine learning and confirmation by immunohistochemistry in colorectal cancer tissue microarrays. Cancers. 2022 May 27;14(11):2664. 10.3390/cancers14112664

154. Kirouac DC, Du J, Lahdenranta J, Onsum MD, Nielsen UB, Schoeberl B, et al. HER2+ cancer cell dependence on PI3K vs. MAPK signaling axes is determined by expression of EGFR, ERBB3 and CDKN1B. Karchin R, editor. PLOS Computational Biology. 2016 Apr 1;12(4):e1004827. 10.1371/journal.pcbi.1004827

155. Arteaga CL, Engelman JA. ERBB receptors: from oncogene discovery to basic science to mechanism-based cancer therapeutics. Cancer cell. 2014 Mar 17;25(3):282–303. 10.1016/j.ccr.2014.02.025

156. Paplomata E, O’Regan R. The PI3K/AKT/mTOR pathway in breast cancer: targets, trials and biomarkers. Therapeutic Advances in Medical Oncology. 2014 Apr;6(4):154–66. 10.1177/1758834014530023

157. Li H, Prever L, Hirsch E, Gulluni F. Targeting PI3K/AKT/mTOR signaling pathway in breast cancer. Cancers. 2021 Jul 14;13(14):3517. 10.3390/cancers13143517

158. Shi P, Feng J, Chen C. Hippo pathway in mammary gland development and breast cancer. Acta Biochimica et Biophysica Sinica. 2014 Dec 2;47(1):53–9. 10.1093/abbs/gmu114

159. Han Y. Analysis of the role of the Hippo pathway in cancer. Journal of Translational Medicine. 2019 Apr 8;17(1). 10.1186/s12967-019-1869-4

160. Zhao M, Mishra L, Deng CX. The role of TGF-β/SMAD4 signaling in cancer. International Journal of Biological Sciences. 2018;14(2):111–23. 10.7150/ijbs.23230

161. Labibi B, Bashkurov M, Wrana JL, Attisano L. Modeling the control of TGF-β/Smad nuclear accumulation by the Hippo pathway effectors, Taz/Yap. iScience. 2020 Aug;23(8). 10.1016/j.isci.2020.101416

162. Band AM, Laiho M. Crosstalk of TGF-β and estrogen receptor signaling in breast cancer. Journal of Mammary Gland Biology and Neoplasia. 2011 Mar 11;16(2):109–15. 10.1007/s10911-011-9203-7

163. Moses H, Barcellos-Hoff MH. TGF-β biology in mammary development and breast cancer. Cold Spring Harbor Perspectives in Biology. 2010 Sep 1;3(1). 10.1101/cshperspect.a003277

164. Zhang Y, Alexander PB, Wang XF. TGF-β family signaling in the control of cell proliferation and survival. Cold Spring Harbor Perspectives in Biology. 2016 Dec 5;9(4). 10.1101/cshperspect.a022145

165. Yang Z, Liu Z. The emerging role of microRNAs in breast cancer. Journal of Oncology. 2020 Jul 3;2020:1–7. 10.1155/2020/9160905

166. Ye L, Wang F, Wang J, Wu H, Yang H, Yang Z, et al. Role and mechanism of miR-211 in human cancer. Journal of Cancer. 2022;13(9):2933–44. 10.7150/jca.71401

167. Shirvani H, Ghanavi J, Aliabadi A, Mousavinasab F, Talebi M, Majidpoor J, et al. Retraction notice to “MiR-211 plays a dual role in cancer development: From tumor suppressor to tumor enhancer”. Cellular Signalling. 2024 Jul 1;121:111266–6. 10.1016/j.cellsig.2024.111266

168. Dabrowski MJ, Wojtas B. Global DNA methylation patterns in human gliomas and their interplay with other epigenetic modifications. International Journal of Molecular Sciences. 2019 Jul 15;20(14):3478. 10.3390/ijms20143478

169. Líu H, Tsai HW, Yang M, Li G, Bian Q, Ding G, et al. Three-dimensional genome structure and function. MedComm. 2023 Jul 8;4(4). 10.1002/mco2.326

170. Calderon L, Weiss FD, Beagan JA, Oliveira MS, Georgieva R, Wang YF, et al. Cohesin-dependence of neuronal gene expression relates to chromatin loop length. eLife. 2022 Apr 26;11. 10.7554/elife.76539

171. Bateman JR, Johnson JE. Altering enhancer–promoter linear distance impacts promoter competition in cis and in trans. Genetics. 2022 Jun 24;222(1). 10.1093/genetics/iyac098

172. Zheng H, Xie W. The role of 3D genome organization in development and cell differentiation. Nature Reviews Molecular Cell Biology. 2019 Jun 13;20:535–50. 10.1038/s41580-019-0132-4

173. Sehgal P, Chaturvedi P. Chromatin and cancer: implications of disrupted chromatin organization in tumorigenesis and its diversification. Cancers. 2023 Jan 11;15(2):466. 10.3390/cancers15020466

174. Bompadre O, Andrey G. Chromatin topology in development and disease. Current opinion in genetics & development. 2019 Apr 1;55:32–8. 10.1016/j.gde.2019.04.007

175. Stephens AD. Chromatin rigidity provides mechanical and genome protection. Mutation Research. 2020 May;821:111712. 10.1016/j.mrfmmm.2020.111712

176. Fischer T, Hayn A, Mierke CT. Effect of Nuclear Stiffness on Cell Mechanics and Migration of Human Breast Cancer Cells. Frontiers in Cell and Developmental Biology. 2020 May 29;8. 10.3389/fcell.2020.00393

177. Meghani K, Folgosa Cooley L, Piunti A, Meeks JJ. Role of chromatin modifying complexes and therapeutic opportunities in bladder cancer. Bladder Cancer. 2022 Feb 28;8(2):101–12. 10.3233/blc-211609

178. Yun JW, Song SH, Kim HP, Han SW, Yi EC, Kim TY. Dynamic cohesin-mediated chromatin architecture controls epithelial–mesenchymal plasticity in cancer. EMBO Reports. 2016 Jul 27;17(9):1343–59. 10.15252/embr.201541852

179. Hao Y, Baker D, ten Dijke P. TGF-β-mediated epithelial-mesenchymal transition and cancer metastasis. International Journal of Molecular Sciences. 2019 Jun 5;20(11):2767. 10.3390/ijms20112767

