## Supplementary material for "Unveiling epigenetic regulatory elements associated with breast cancer development": S1_Fig

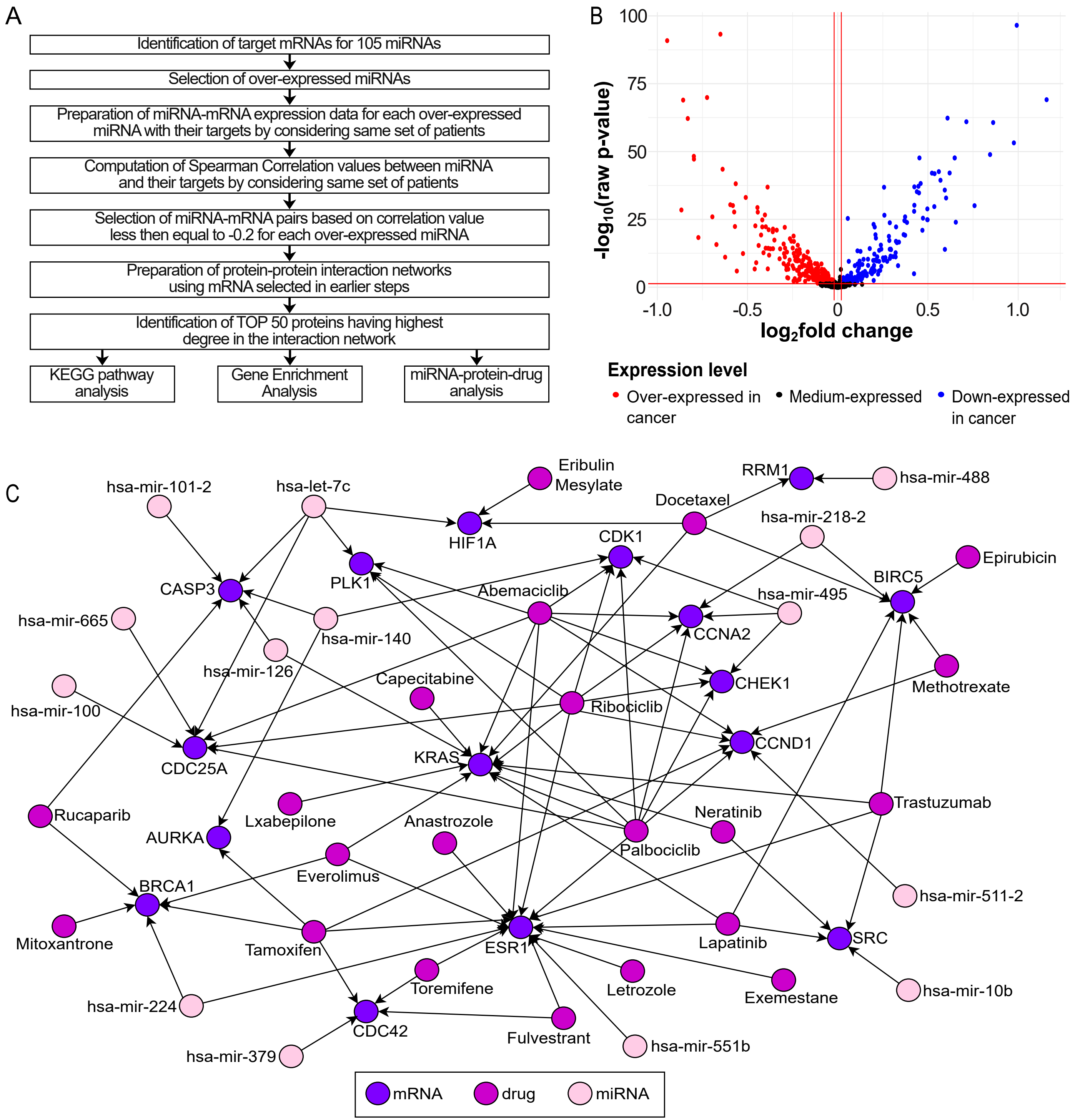


**S1 Fig** Suppressive impact of putative miRNA markers. (A) Pipeline to identify suppressive impact of miRNA on mRNA. (B) Volcano plot of miRNA genes expression. (C) miRNA-protein-drug network.
