## Supplementary material for "Unveiling epigenetic regulatory elements associated with breast cancer development": S2_Fig

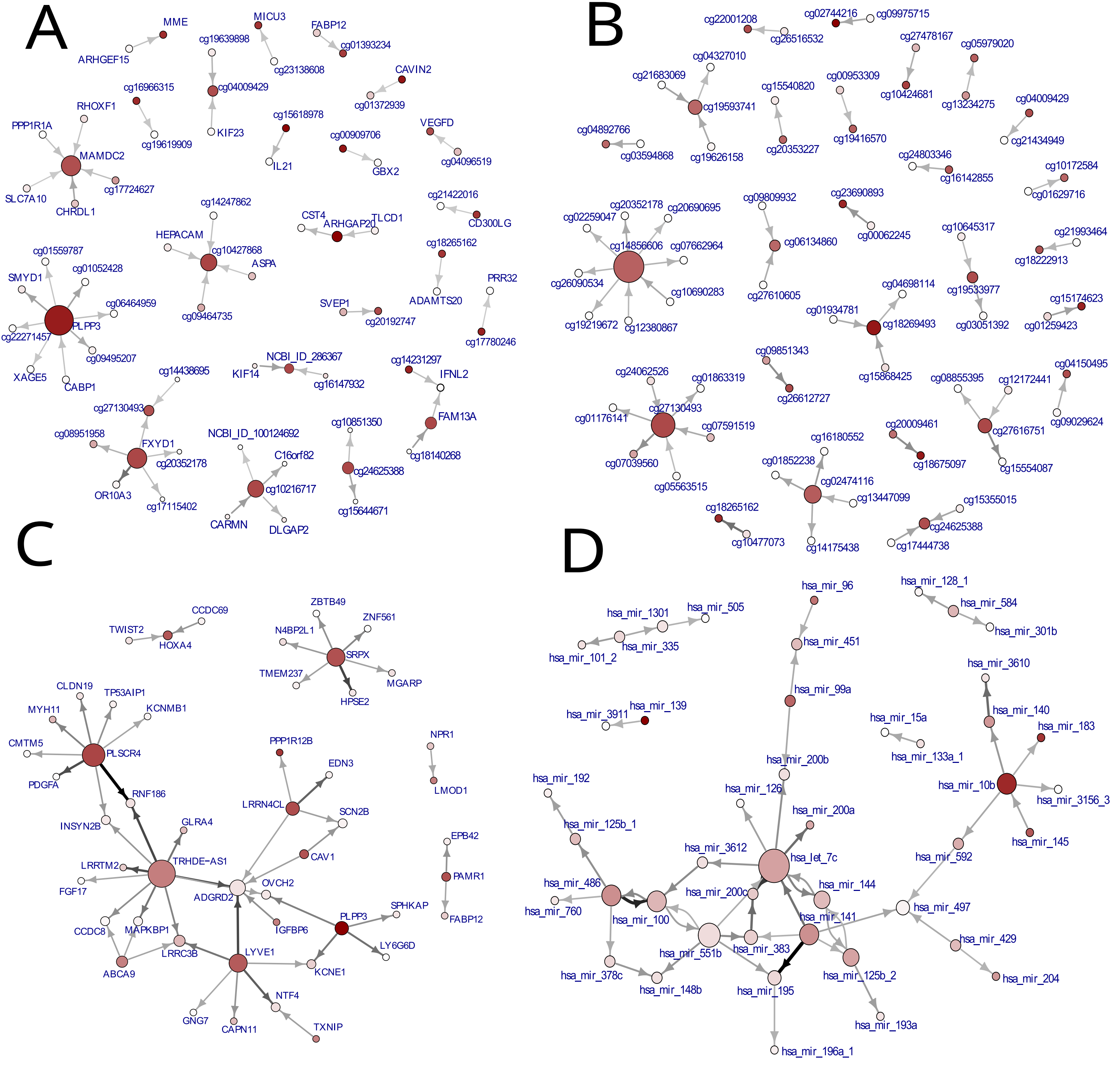


**Fig S2** Interaction graphs (ID-Graphs) obtained from MCFS-ID. Interactions between significant features in the context of classification cancer/normal patients. (A) all categories (B) single category: DNA methylation data (C) single category: mRNA data (D) single category: miRNA data
