## Supplementary material for "Unveiling epigenetic regulatory elements associated with breast cancer development": S3_Fig

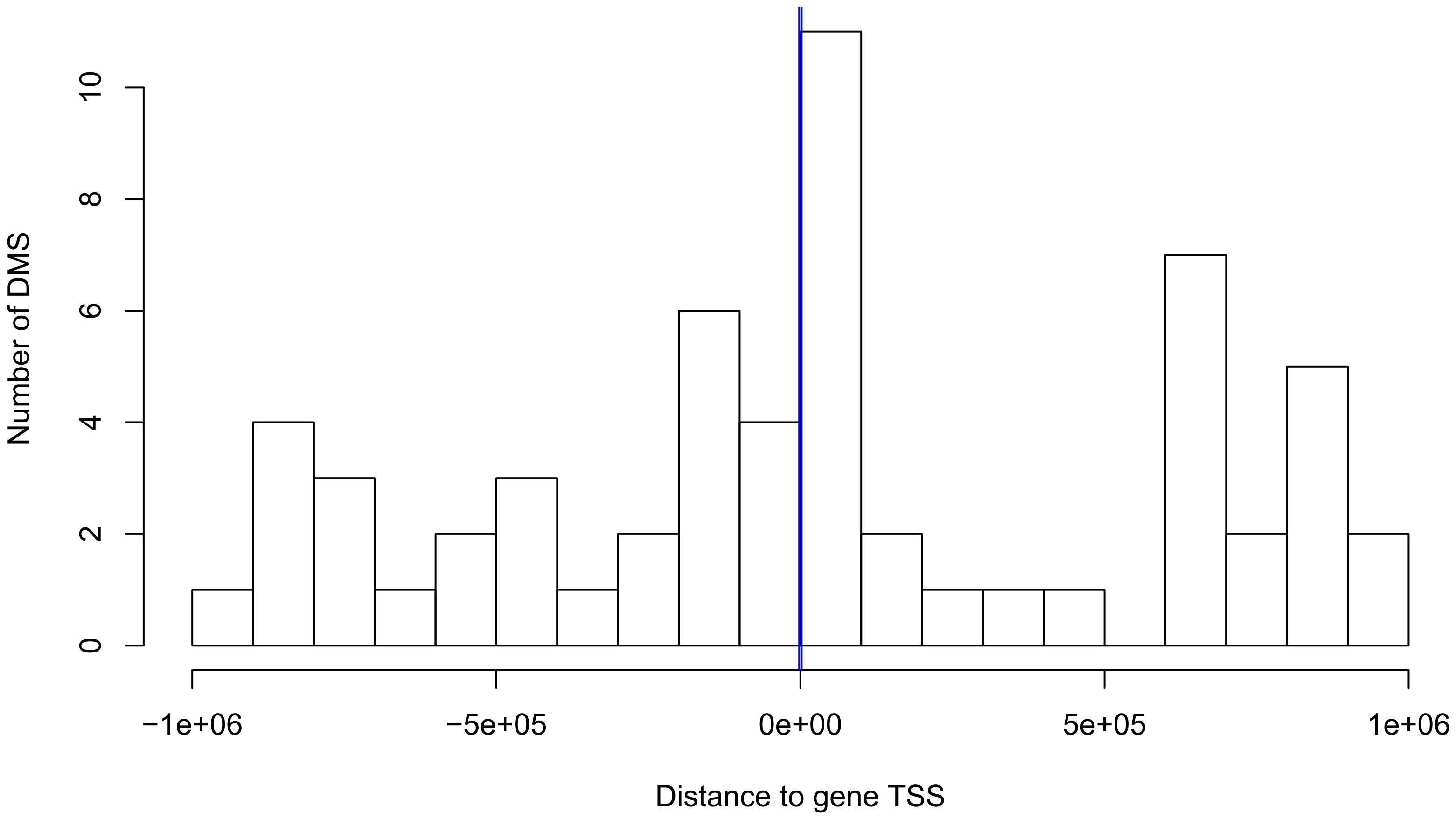


**S3 Fig** Distribution of distances between gene TSS and DMS that were found to correlate significantly (Spearman correlation, FDR<0.05, |rho| ≥0.6). Correlation was performed between gene expression and DMSs located 1Mbp from a gene TSS. The blue line indicated the promoter region ± 2000bp from a TSS.
