## Supplementary material for "Unveiling epigenetic regulatory elements associated with breast cancer development": S4_Fig

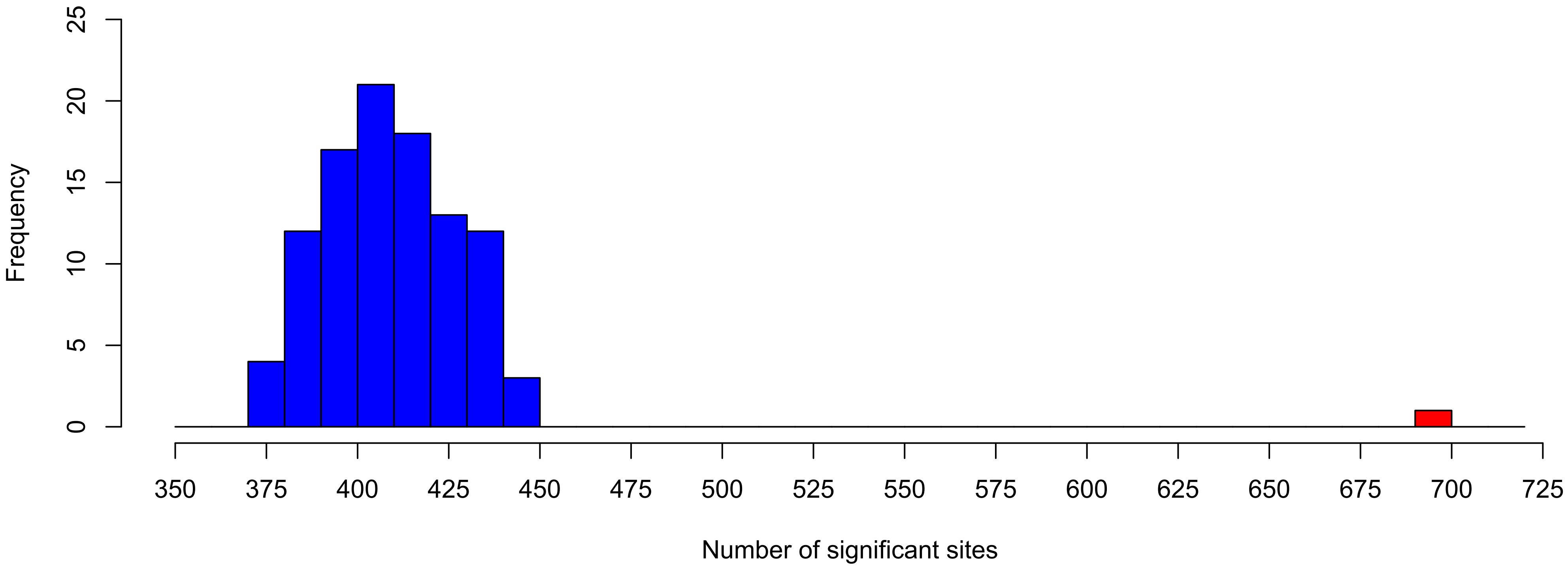


**S4 Fig** Distribution of the number of sites that had a significant impact on the patient survival. Blue - distribution of significant sites from 2006 sites selected randomly using a bootstrapping technique (sampling 100 times); red - number of significant sites from a set of DMS.
