## Supplementary material for "Unveiling epigenetic regulatory elements associated with breast cancer development": S5_Fig

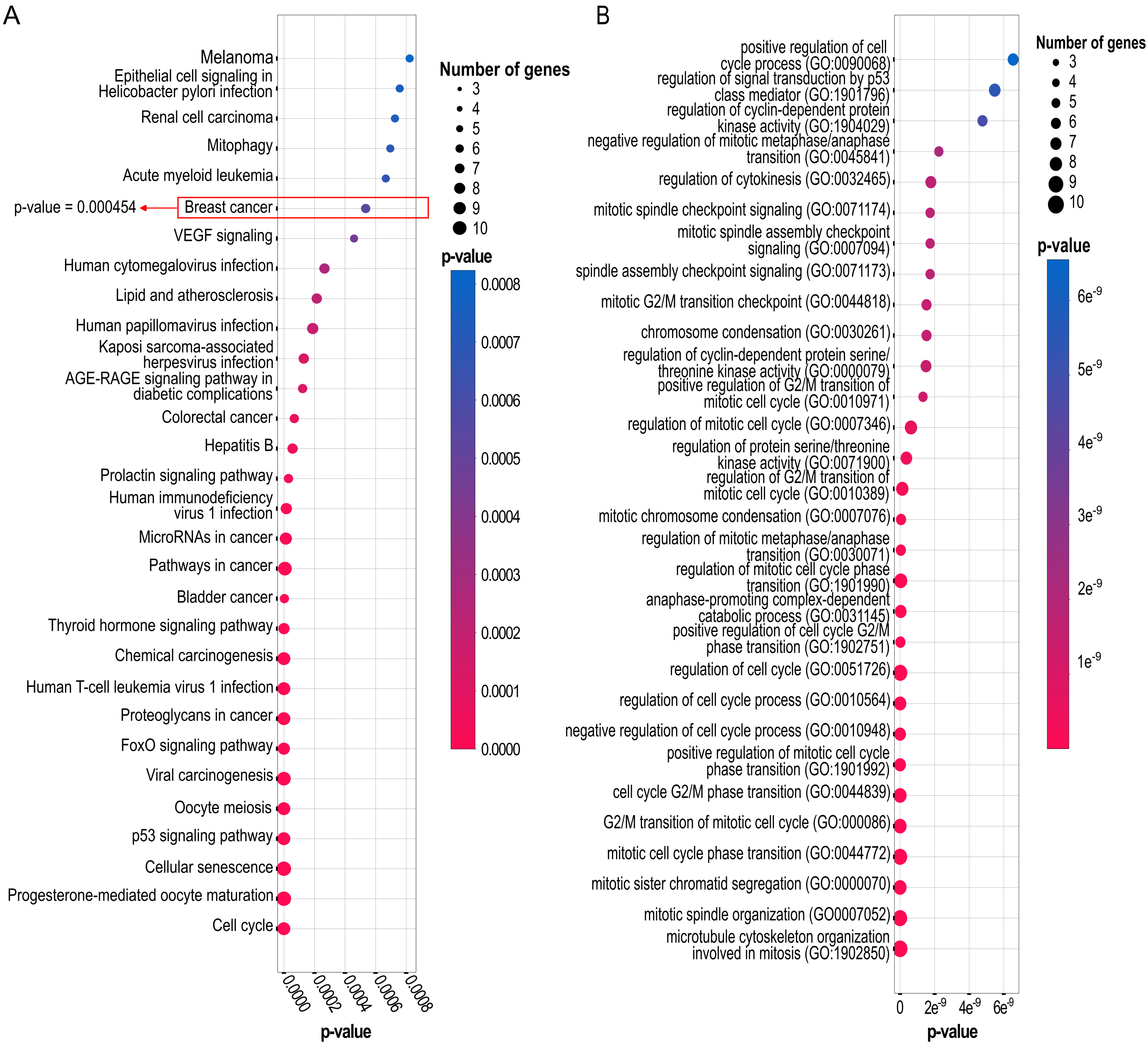


**S5 Fig** Functional analysis of the over-expressed genes encoding 50 proteins with the highest number of interaction network (A) KEGG pathway analysis (B) GO BP analysis
