## Supplementary material for "Unveiling epigenetic regulatory elements associated with breast cancer development": S6_Fig

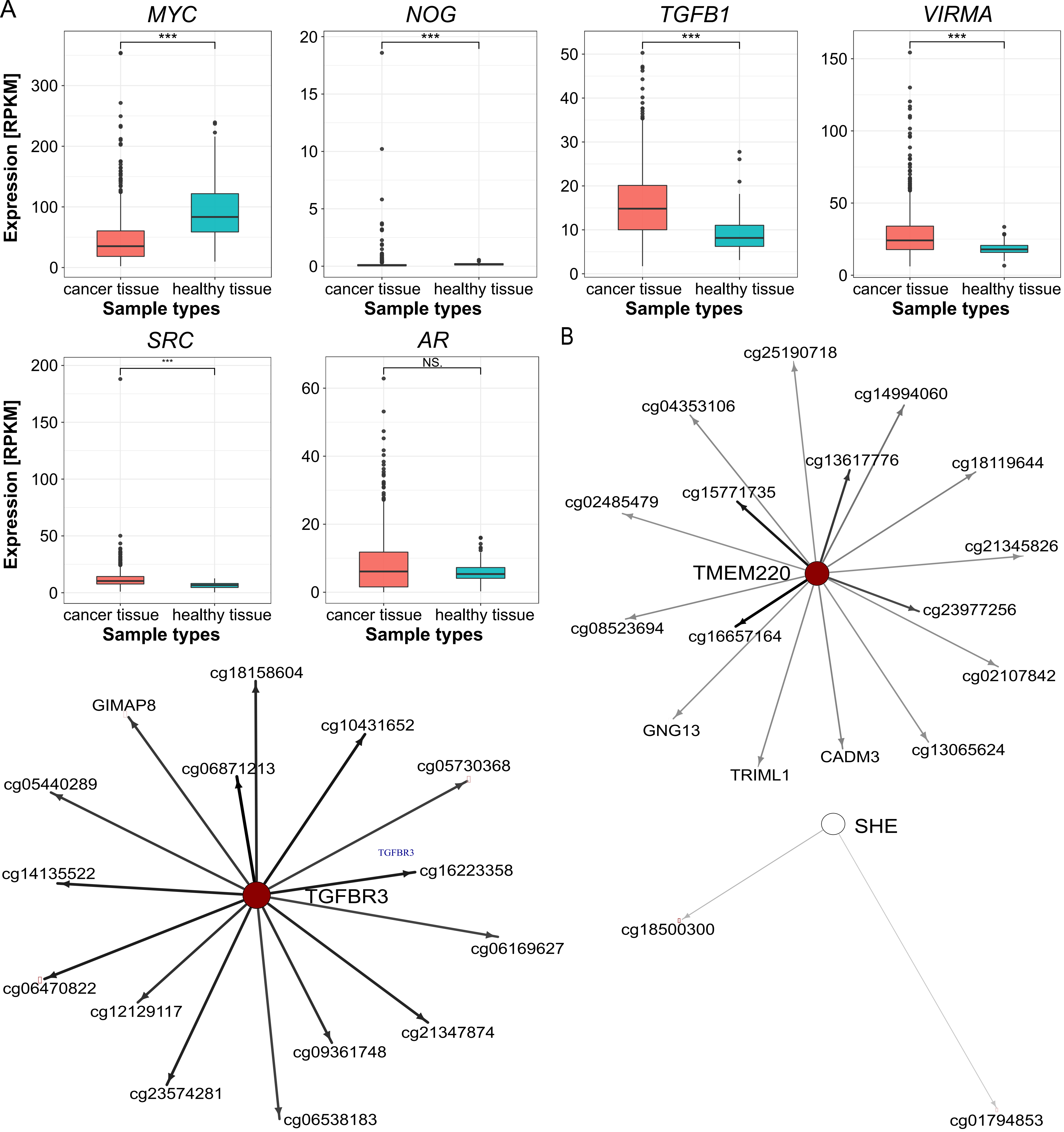


**S6 Fig** Additional profiles for genes in regulatory networks (A) Expression profiles of genes with the largest number of putative interactions in gene-gene interactions network. Wilcoxon test was used to compare significance of expression changes between them and *** means p-value≤0.001. (B) Interaction graphs (ID-Graphs) obtained from MCFS-ID for three target genes whose linear models reached statistical significance p≤0.05 and R2>0.5 [*TMEM220, TGFBR3, SHE*].
