## Supplementary material for "Unveiling epigenetic regulatory elements associated with breast cancer development": S7_Fig

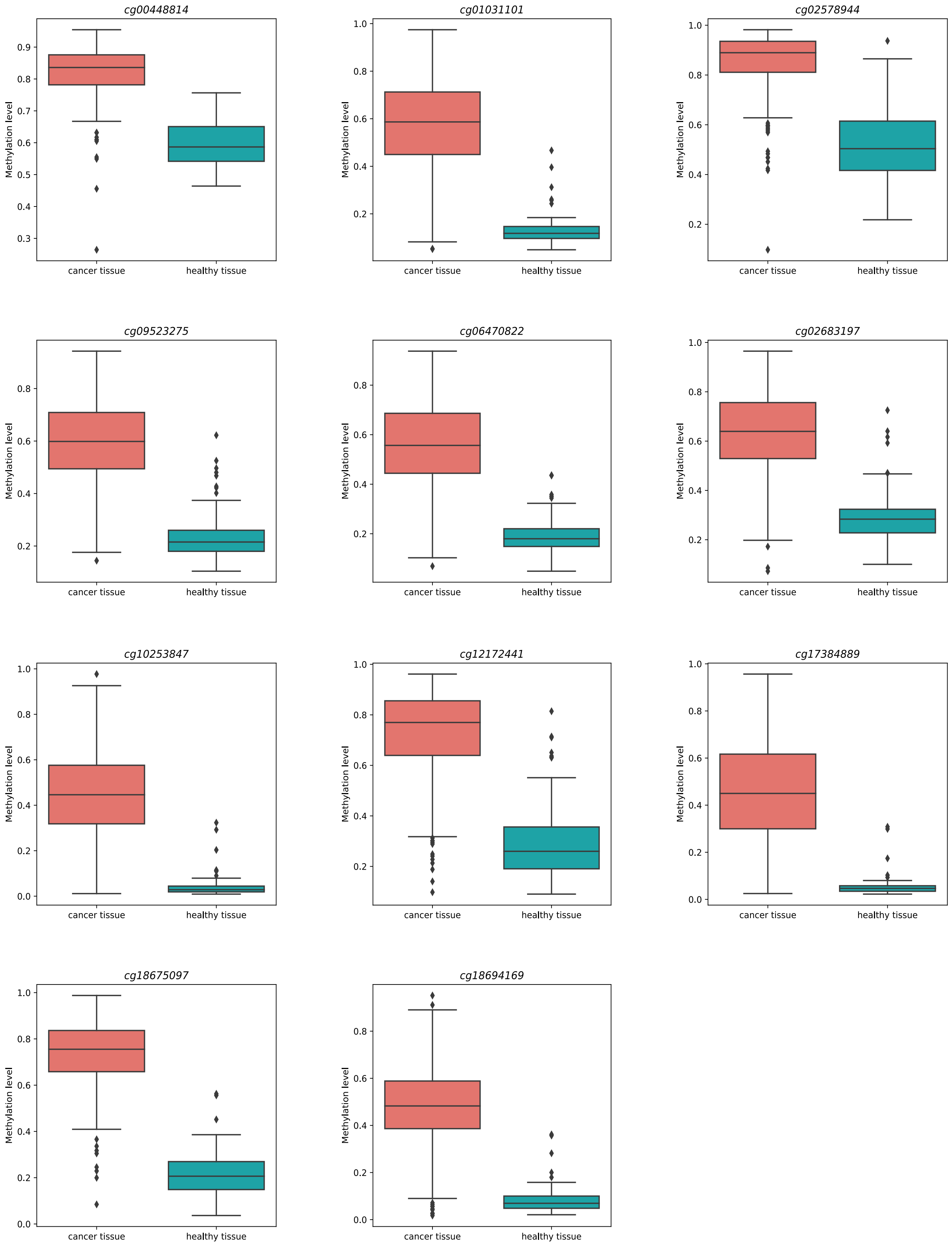


**S7 Fig** Methylation profile of significant methylation around the *NKAPL* gene for cancer tissue (MCF-7) in red and healthy tissue (MCF-10A) in blue.
